# Disagreement among variant effect predictors guides experimental prioritization of target proteins

**DOI:** 10.64898/2026.03.18.712765

**Authors:** Nicolas F. Jonsson, Joseph A. Marsh, Kresten Lindorff-Larsen

## Abstract

Interpreting the functional consequences of genetic variation, especially rare missense variants, remains a significant challenge in human genetics. Computational variant effect predictors (VEPs) and multiplexed assays of variant effects (MAVEs) provide complementary approaches, with VEPs offering scalable predictions and MAVEs delivering detailed empirical measurements. However, MAVEs are resource intensive and cannot yet be applied broadly across the proteome, making it important to identify proteins where experimental mapping will be most informative. We hypothesised that MAVEs should be particularly valuable for proteins where computational predictors disagree, as such disagreement may highlight mechanistic blind spots. To test this, we analysed predictions from ten distinct VEPs across more than 13,000 human proteins and quantified inter-predictor concordance. We observed substantial variability across proteins in the degree of agreement across predictors and investigated structural, functional and gene-level features associated with this variation. We find that inter-VEP concordance showed no relationship with agreement to experimental MAVE data. If predictor agreement reflected how intrinsically predictable a protein is, these quantities would be expected to correlate. Their decoupling instead suggests that MAVEs may provide orthogonal information to VEPs, supporting the use of inter-VEP disagreement to prioritise proteins where experimental data will be most informative. We therefore propose using inter-VEP disagreement as a practical strategy to prioritise proteins for experimental characterization. Focusing on proteins with low predictor concordance should maximise the informational value of new MAVEs, and improve variant interpretation in both research and clinical contexts.

## Introduction

Understanding the consequences of genetic variants is critical for unravelling the molecular mechanisms underlying human health and disease. Variant effect predictors (VEPs) have become central computational tools for estimating how genetic variants, particularly missense variants, influence protein function, structure, and stability (***Correa Marrero et al., 2024***; ***Horne and Shukla, 2022***; ***Fawzy and Marsh, 2024***; ***Ng and Henikoff, 2003***; ***Livesey and Marsh, 2022***). By facilitating large-scale interpretation of human genetic variation, VEPs play a pivotal role not only in fundamental biological research but also in clinical diagnostics, supporting not only experimental validations but also clinical interpretation of genetic disorders (***Telenti et al., 2018***; ***Brandes et al., 2023***; ***Pejaver et al., 2022***; ***Livesey and Marsh, 2022***).

Historically, early predictive approaches relied on methods such as substitution matrices (e.g., BLOSUM62) (***Henikoff and Henikoff, 1992***) or heuristic models informed by evolutionary conservation, amino acid properties, and structural insights. Tools like SIFT (***Ng and Henikoff, 2003***) and PolyPhen-2 (***Adzhubei et al., 2010***) demonstrated that combining multiple sequence alignments with physicochemical and structural heuristics could effectively differentiate likely pathogenic from benign variants, establishing foundational principles for subsequent advancements (***Fawzy and Marsh, 2025***; ***Livesey and Marsh, 2020***). Over the past two decades, advances in machine learning techniques and expanded biological datasets have revolutionised the VEP landscape, increasing the complexity and depth of predictive models, leading to more sophisticated predictors that integrate evolutionary, biophysical, and structural features, often through machine learning and deep learning frameworks. Contemporary VEPs integrate extensive biophysical-, evolutionary-, and structural features, frequently employing deep neural networks and protein language models trained on massive protein datasets (***Meier et al., 2021***; ***Rives et al., 2021***; ***Hsu et al., 2022***; ***Horne and Shukla, 2022***; ***Ofer et al., 2021***; ***Hu et al., 2024***). Notably, some of these predictors are trained on clinically annotated variants and are often further tuned using patterns of population variation, which can shape their learned decision boundaries. Additionally, the recent advent of highly accurate protein structure predictions, exemplified by AlphaFold2 (***Jumper et al., 2021***), has significantly broadened available feature sets, allowing coverage of proteins lacking experimentally solved structures to be assessed with detailed structural context (***Gerasimavicius et al., 2025***; ***Tran et al., 2024***; ***Telenti et al., 2018***; ***Correa Marrero et al., 2024***; ***Cagiada et al., 2025***). Despite these major advances, predictive accuracy still remains uneven across different proteins and contexts. Sensitivity and specificity are often limited where functional effects are subtle or evolutionary signal is weak—most notably in intrinsically disordered (IDRs) and other low-conservation regions, where conformational flexibility obscures constraints (***Fawzy and Marsh, 2025***; ***Uversky et al., 2008***; ***Chow et al., 2024***; ***Cagiada et al., 2025***). In such settings, models may even display high mutual agreement driven by shared inductive biases (e.g., conservation metrics or allele frequency priors) rather than correctness—a ‘false consensus’. At the same time, VEP performance and inter-predictor concordance are highly heterogeneous across proteins and regions, with particularly poor behaviour in intrinsically disordered and other structurally complex contexts. Prior work has shown that many clinically relevant variants in these regions map to short linear motifs, interaction interfaces, and regulatory elements that are mechanistically complex and difficult for current feature sets to model (***Fawzy and Marsh, 2024***, ***2025***; ***Livesey and Marsh, 2025***; ***Alderson et al., 2023***). Together, these observations motivate the hypothesis that inter-VEP disagreement may preferentially arise in biologically complex contexts and may therefore highlight candidate blind spots for empirical investigation.

In parallel with the computational developments, multiplexed assays of variant effects (MAVEs) have matured into powerful, high-throughput experimental platforms capable of mapping the functional landscape of thousands of variants at once (***Findlay et al., 2018***; ***Weile et al., 2017***; ***Starita et al., 2017***). These assays generate high-resolution maps of variant impacts, providing independent empirical benchmarks free from many biases in clinical training data. Additionally, MAVEs have revealed strong correspondence between functional and clinical classifications in some well-characterised domains, and have helped assign functional status to many variants (***Findlay et al., 2018***; ***Dace et al., 2025***; ***Livesey and Marsh, 2025***). However, despite their power, MAVEs are resource-intensive and technically demanding, leaving substantial experimental gaps—particularly for complex, multifunctional proteins—and thus cannot yet be applied exhaustively across the human proteome (***Zheng et al., 2023***), making strategic prioritization essential. Given the impracticality of exhaustively testing all proteins experimentally, determining optimal allocation of limited experimental resources has become increasingly critical. While VEPs can rapidly generate scalable predictions, discrepancies between VEPs may indicate problematic regions for computational methods or highlight biological nuances distinct from those captured by experimental assays. However, whether consensus among different VEPs reliably reflects functional ground truth as measured by MAVEs, or whether regions of VEP disagreement represent important predictive ‘blind spots’ warranting experimental follow-up, remains uncertain.

In this study, we evaluate the concordance of variant effect predictions across ten state-of-the-art VEPs, covering over 13,000 human proteins. We quantify global and context-specific inter-VEP agreement, investigating how features such as protein type, gene-level characteristics, and population genetic annotations may explain prediction discrepancies. We assess whether inter-VEP consensus corresponds to empirical functional measurements provided by MAVEs, testing if computational agreement can indeed serve as a proxy for functional accuracy. Our findings highlight substantial variability in inter-VEP concordance across the human proteome and demonstrate no correlation between VEP consensus and MAVE-based empirical results. Consequently, we propose using areas of inter-VEP disagreement as strategic targets for empirical investigation, thereby optimizing experimental resource allocation, enhancing future VEP benchmarking, and refining variant interpretation in both research and clinical practice.

## Results and Discussion

### Methodological lineages shape concordance among VEPs

To characterise the reliability and variability of current VEPs, we began by assembling a large dataset of missense variant predictions (Figure 1). Our dataset was built from publicly available resources, incorporating outputs from 71 distinct VEPs, mirroring the scope and methodology of prior large-scale efforts (***Livesey and Marsh, 2025***). These models vary widely in both their input features and training paradigms, reflecting the full breadth of the contemporary prediction landscape. Prior to making the comparisons, we polarity-aligned the predictor scores so that greater values consistently corresponded to predictions of more deleterious variants, following the authors’ documentation or, when ambiguous, the polarity that maximised agreement with benchmark multiplexed assays (see Methods). To enable direct comparison across predictors, we constructed a harmonised dataset of single-amino acid substitutions accessible via single-nucleotide variants (SNVs) that were jointly scored by all VEPs. This shared dataset was essential for eliminating coverage-related biases, ensuring that observed differences in prediction were attributable to model behaviour rather than discrepancies in input space (Fig. 2). To assess whether restricting the analysis to universally scored variants affected the inferred similarity landscape, we also performed a pairwise analysis of VEP outputs to maximise the number of variants considered in each comparison (Fig. S1). To verify that restricting the analysis to universally scored variants did not distort the inferred similarity landscape, we repeated the analysis using the maximally over-lapping set of variants jointly scored by each predictor pair, including both SNV-accessible substitutions and multi-nucleotide variants (MNVs) where available. The resulting correlation structure closely matched that obtained from the globally shared SNV dataset (*R*^2^ = 0.959, see Supplementary Fig. S1), indicating that the observed concordance patterns are robust to the choice of variant inclusion strategy. For consistency, interpretability, and to minimise variation in input composition across comparisons, we therefore used the all-shared SNV dataset as the basis for the remainder of our analyses.

**Figure 1.**
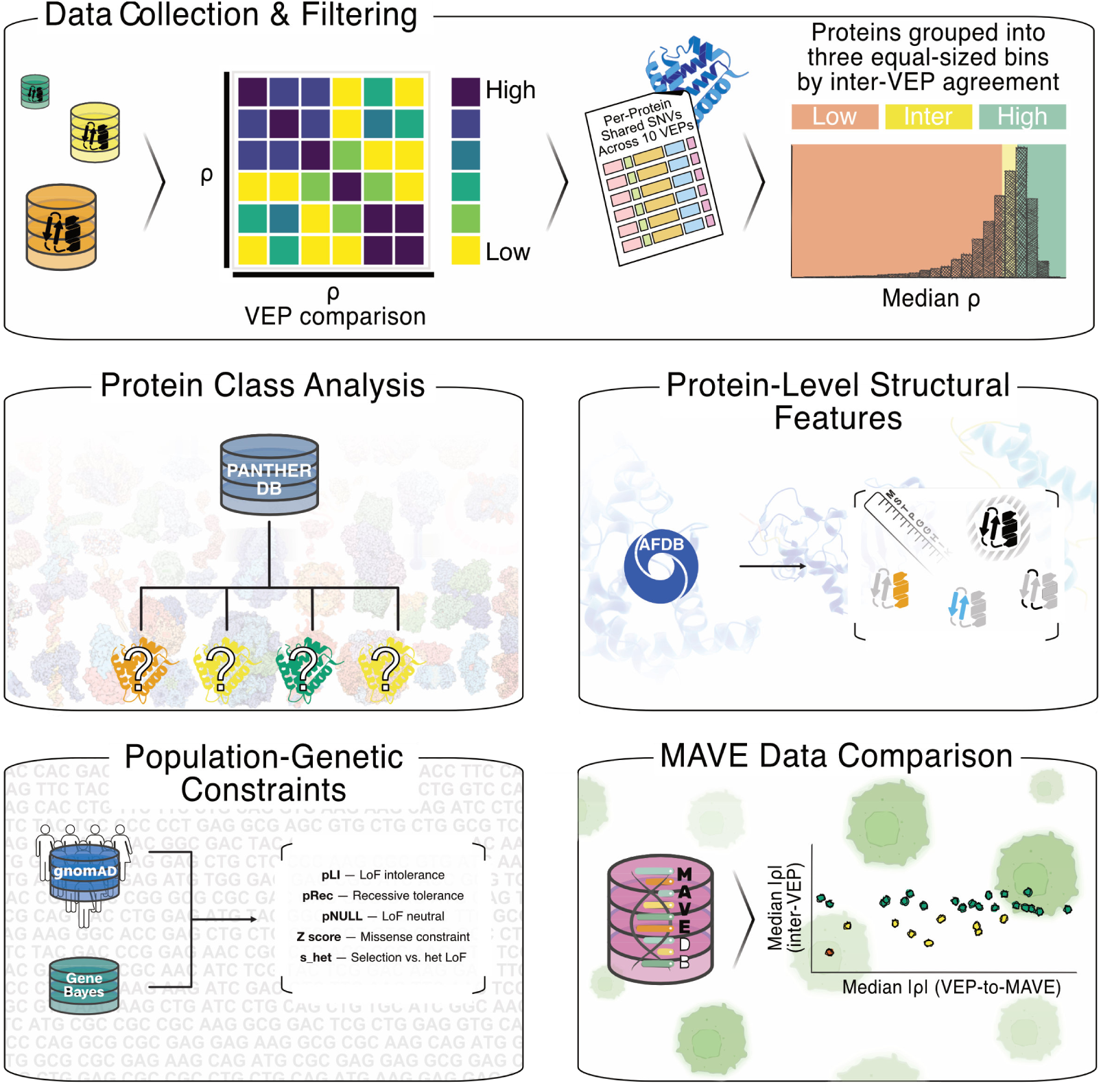
Graphical summary of our characterisation of VEP limitations across multiple levels, from model comparison to target nomination. Tiles (left-to-right, top-to-bottom) depict: (i) assembly of predictions from 71 VEPs and construction of a harmonised variant set; (ii) global concordance mapping via Spearman correlations and hierarchical clustering; (iii) per-protein concordance and tiering into low, intermediate, and high agreement; (iv) class-level trends using PANTHER annotations; (v) structure-informed features from AlphaFold2 (pLDDT, rASA/WCN, secondary structure) summarised per protein; (vi) gene-level population-genetic constraints from gnomAD (pLI, pRec, pNull, missense Z, *S*_het_); (vii) benchmarking against multiplexed assays of variant effects (MAVEs). Colours follow the agreement tiers used throughout (low, intermediate, high). Data is available in an online database (see Code and Data Availability).

**Figure 2.**
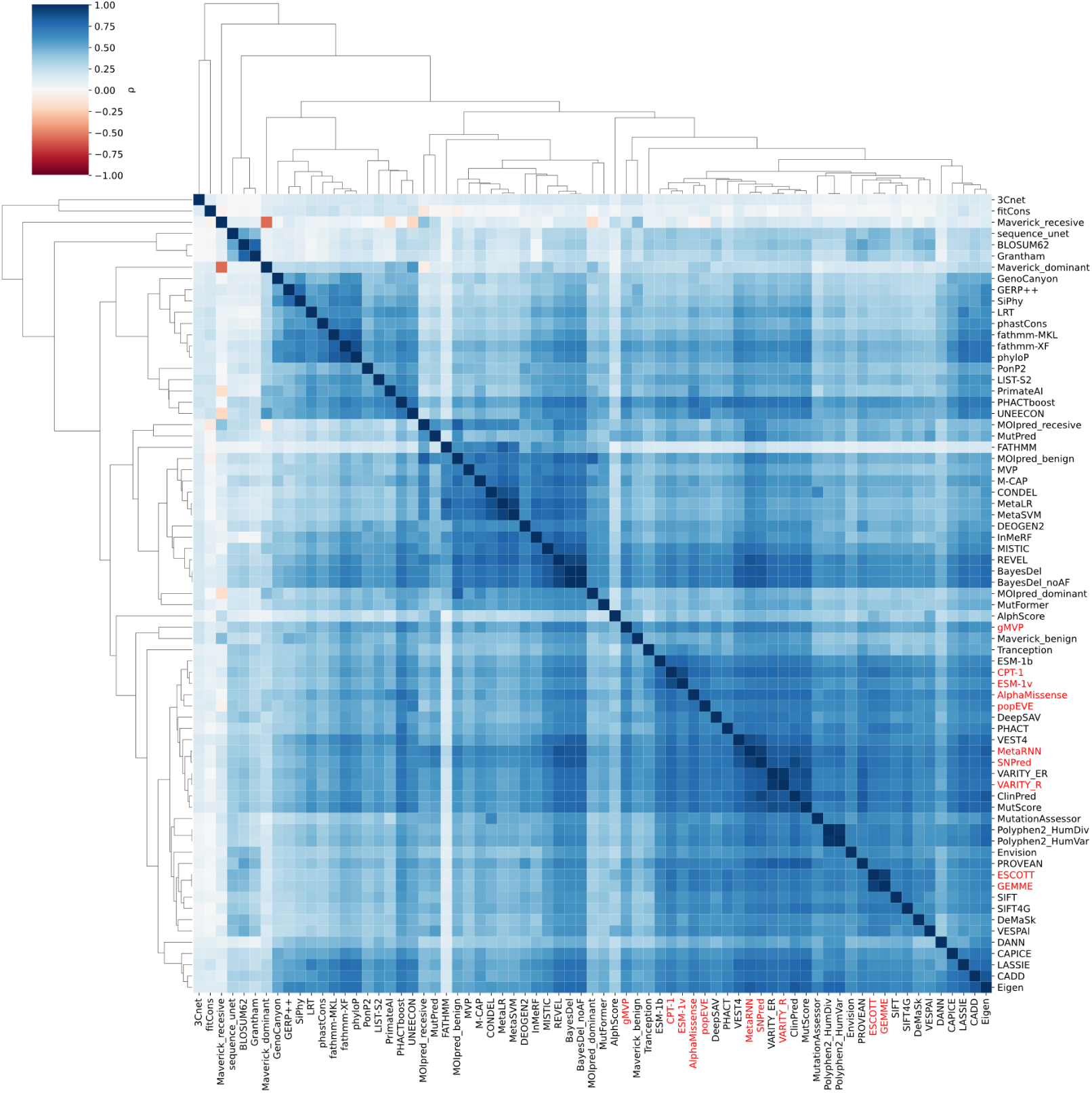
Global concordance landscape across 71 VEPs. The heatmap shows Spearman correlations between predictors computed on a harmonised set of single-nucleotide variants jointly scored by all methods. Predictor outputs were first polarity-aligned so that higher scores consistently indicate greater predicted deleteriousness. Rows and columns are ordered by hierarchical clustering of the correlation profiles using correlation distance and average linkage, grouping predictors with similar patterns of agreement across the full panel of methods. Distinct blocks correspond to families of related predictors, including embedding and protein language-model approaches (e.g., AlphaMissense, CPT-1, popEVE, ESM-1b/1v), alignment-driven methods (GEMME, ESCOTT), and clinically trained or ensemble predictors (MetaRNN, SNPred, VARITY_R). The ten predictors used in downstream analyses are highlighted in red on the axes. The colour scale represents Spearman correlation coefficients, where positive values indicate concordant variant ranking between predictors, values near zero indicate little rank agreement, and negative values indicate opposing ranking tendencies. Analyses using pairwise-overlapping variant sets for each predictor pair yield the same qualitative structure (Supplementary Fig. S1).

Inspection of the predictor relationships revealed structured clustering among models, not surprisingly reflecting shared methodological foundations. This is illustrated in the clustermap of Spearman correlations across all 71 VEPs, calculated over the dataset of shared SNVs after polarity alignment so that higher scores consistently indicate greater predicted deleteriousness (Fig. 2). Several densely connected blocks emerge along the diagonal, indicating that certain groups of predictors consistently exhibit stronger mutual correlation than others. These clusters correspond to known design similarities. For instance, AlphaMissense, CPT-1, popEVE, and ESM-1v group together with other embedding-based models, reflecting their common use of large-scale protein language models and unsupervised sequence representations. GEMME and ESCOTT form a separate cluster aligned with their empirical, alignment-driven methodologies that rely on multiple sequence alignments and conservation metrics. Meanwhile, clinically trained predictors such as MetaRNN, SNPred, and gMVP appear more isolated, showing only moderate correlation with each other and reduced similarity to the rest of the predictor set.

From this unified dataset, we curated a representative subset of ten VEPs for in-depth concordance analysis. This subset was chosen to strike a balance between empirical performance, proteome coverage, and methodological diversity. First, we prioritised predictors with strong experimental accuracy, based on benchmarks against data generated by MAVEs (***Livesey and Marsh, 2023***). In parallel, we assessed internal data coverage at the protein level, favouring models that scored a higher proportion of residues within individual proteins. We also considered proteome-wide availability to ensure a sufficiently broad set of proteins for generalizable comparisons. Finally, to avoid over-representation of any single modelling approach, we selected VEPs spanning a range of training philosophies (***Pathak et al., 2024***; ***Livesey and Marsh, 2025***), including population-agnostic models, population-tuned predictors that incorporate allele frequency data, and clinically trained tools developed using labelled disease variants. This selection was intended not only to facilitate robust concordance analysis but also to explore how differing algorithmic lineages influence prediction agreement. Importantly, the ten VEPs selected for focused analysis—highlighted in red in Fig. 2—are distributed across these distinct regions, reinforcing that while they demonstrate broad global agreement, they also capture diverse aspects of variant impact. This pattern highlights that predictor disagreement is not random but rather shaped by differences in input features, training data, and biological factors, the latter a point that we investigate further in the following sections. The final panel of predictors included AlphaMissense (***Cheng et al., 2023***), CPT-1 (***Jagota et al., 2023***), GEMME (***Laine et al., 2019***), ESCOTT (***Tekpinar et al., 2025***), ESM-1v (***Meier et al., 2021***), popEVE (***Orenbuch et al., 2025***), VARITY_R (***Wu et al., 2021***), MetaRNN (***Li et al., 2022***), SNPred (***Molotkov et al., 2023***), and gMVP (***Zhang et al., 2022***). These VEPs represent a spectrum of methodological strategies, from empirically grounded tools like GEMME and ESCOTT, to deep learning models such as ESM-1v and gMVP, and ensemble methods like VARITY_R and MetaRNN. The set includes predictors trained on a diversity of sources—from protein sequence embeddings and structural features to clinical databases and population variation. AlphaMissense and popEVE exemplify population-tuned models trained on large-scale population genetics resources, while tools such as SNPred and MetaRNN reflect clinical training using disease-labeled data. This diversity in input features, modelling architecture, and training objectives provides a robust foundation for assessing the consistency—and inconsistency—of variant effect predictions across the proteome.

### Inter-VEP concordance varies across protein functional classes

Having established the overall agreement landscape among a broad range of different VEPs and selected a subset of VEPs from which to calculate inter-VEP concordance, we next extended our analysis to characterise how this concordance varies across individual proteins, aiming to identify local contexts and biological features associated with predictor disagreement. For each protein, we quantified concordance between VEPs by calculating Spearman correlations for all model pairs, using SNVs shared across the full set of ten selected predictors after polarity alignment of predictor scores. We used the Spearman’s rank correlation coefficient to accommodate the often non-linear relationships, capturing relative rank agreement rather than direct similarity of raw prediction scores (***Altman, 1999***). We then calculated the median of all pairwise correlation coefficients for each protein, producing a single measure of inter-VEP agreement that could be compared across multiple proteins. This analysis revealed a broad, continuous distribution of concordance scores across 13,332 proteins with median correlation coefficients varying from 0.025 (TMEM269/A0A1B0GVZ9/Transmembrane protein 269) to 0.924 (ERO1A/Q96HE7/ERO1-like protein alpha) (Fig. 3A). To facilitate downstream comparisons, we partitioned the dataset into three equally-sized categories—low-, intermediate-, and high-agreement—based on these concordance scores.

**Figure 3.**
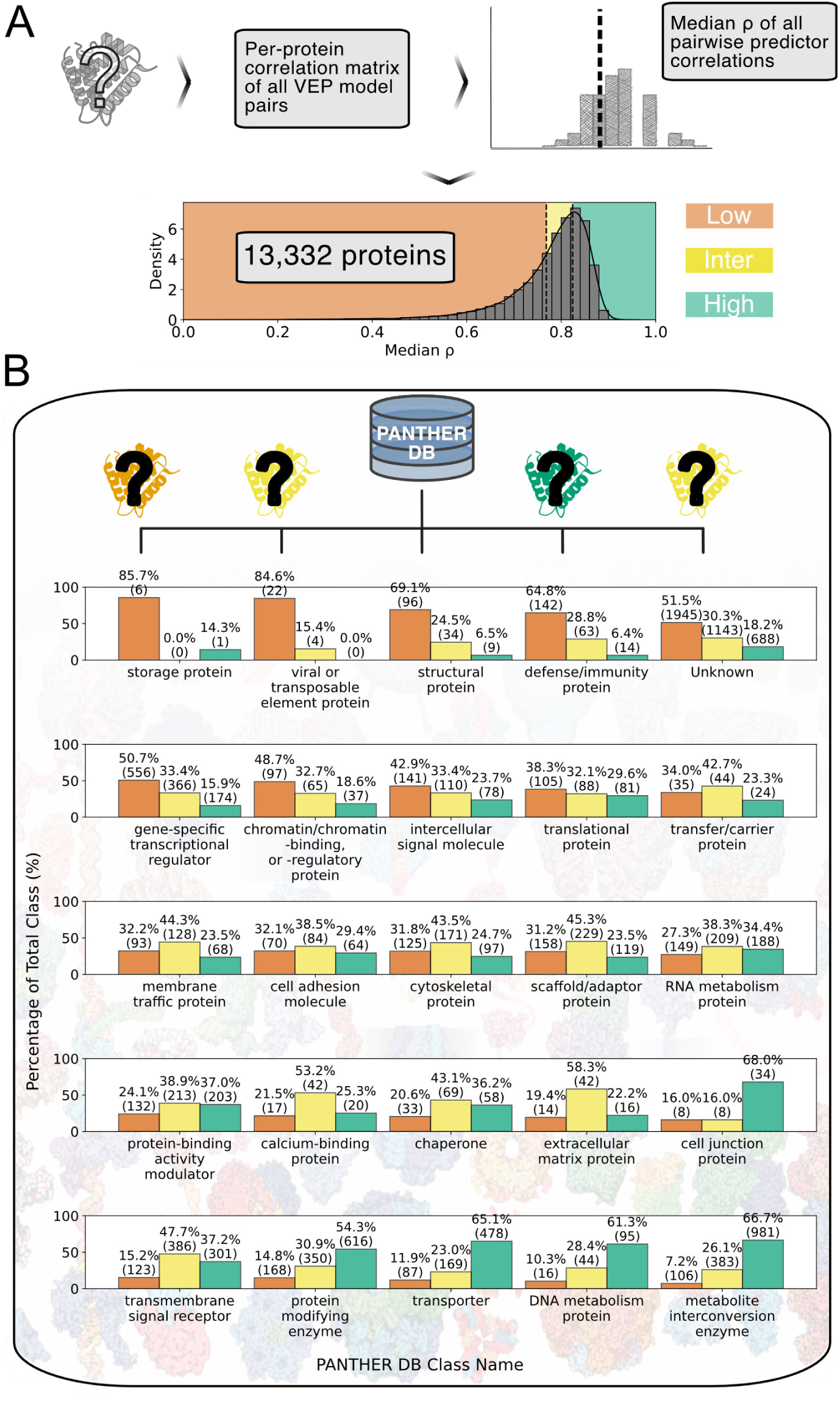
Protein-level variation in inter-VEP agreement. (A) Distribution of per-protein inter-VEP concordance—defined as the median Spearman correlation coefficient across all pairwise combinations of the ten selected VEPs after polarity alignment of predictor scores—computed on the shared-SNV set for *>*13,000 human proteins. The broad, continuous spread motivates the low-, intermediate-, and high-agreement tiers used throughout. (B) Distribution of proteins across top-level PANTHER protein classes, stratified by inter-VEP concordance. Bars show, within each class, the percentage of proteins assigned to low (red), intermediate (yellow), or high (green) agreement based on the per-protein median Spearman correlation. Thresholds match those used in the main analysis, enabling comparison across figures.

First, we investigated whether there were certain functional classes where model disagreement was more pronounced than others. To explore this, we grouped proteins into top-level protein classes based on PANTHER DB database annotations (***Mi et al., 2021***) and assessed the distribution of VEP concordance categories within each class (Fig. 3B). While the overall concordance landscape showed a wide and continuous spread across the proteome, we observed distinct class-specific trends. A subset of protein classes exhibited a notably higher fraction of proteins with low inter-VEP agreement, including storage proteins, viral or transposable element proteins, structural proteins, and defense/immunity proteins.

Storage proteins typically fulfil buffering or nutrient storage functions rather than convey precise molecular interactions (***Recalcati et al., 2008***; ***Broyard and Gaucheron, 2015***). As a result, these proteins tend to be under relaxed purifying selection and show elevated substitution rates, allowing for greater sequence flexibility without compromising function (***Rijnkels, 2002***). Together, these attributes may cause such proteins to lack the evolutionary signatures or structural features that VEPs typically leverage, leading to greater disagreement among models. Similarly, proteins derived from viral or transposable element origins often possess non-canonical sequence characteristics and evolve under atypical constraints or high mutation rates (***Kidwell and Lisch, 2000***; ***Kordišs et al., 1999***; ***Iwasaki et al., 2025***). These proteins can be poorly represented in reference alignments and databases, reducing the accuracy of homology-based and population-trained predictors (***Eddy, 2011***; ***Lawlor and Ellison, 2023***). Consistent with this, recent benchmarks of protein language models (pLMs) on MAVE datasets have shown that viral proteins are predicted substantially less accurately than other proteins, with a simple site-mean baseline often matching or outperforming supervised models (***Vieira et al., 2026***). This behaviour is driven by strong site-specific effects and characteristic variability patterns in viral datasets associated with poor model generalization (***Vieira et al., 2026***). Together, these properties provide a mechanistic explanation for the pronounced inter-VEP disagreement observed in viral-derived protein classes. Structural proteins—such as collagen and actin—play central roles in maintaining cellular and extracellular architecture. Their primary function lies not in discrete biochemical interactions or enzymatic specificity but in maintaining mechanical stability and tissue integrity (***Xu et al., 1998***; ***Letort et al., 2015***). These functions often rely on higher-order quaternary structural assemblies, such as fibrils or filaments, rather than isolated monomeric structures (***Pegoraro et al., 2017***). As a result, the functional consequences of mutations in these proteins may only become apparent in the context of supramolecular complexes, which are more difficult to extract by typical monomer-based or sequence-only variant effect predictors. Defense and immunity proteins, such as antigen receptors, Toll-like receptors (TLRs), and cytokines, are subject to strong diversifying selection due to host-pathogen arms races (***Hughes, 1999***; ***Barreiro and Quintana-Murci, 2010***; ***Minias and Vinkler, 2022***; ***Melepat et al., 2024***; ***Iwasaki et al., 2025***). Domain architectures such as leucine-rich repeats (LRRs) and immunoglobulin folds often display rapid turnover or high variation in specific loops, while maintaining structural scaffolds (***Persi et al., 2016***). This results in species-specific sequence variation and fast-evolving domains, making accurate functional impact prediction particularly challenging for methods that depend on conservation or structural regularity (***Lazzaro and Clark, 2012***; ***Sironi et al., 2015***; ***Ahmad et al., 2021***; ***Markov et al., 2023***). Together, these patterns suggest that VEP disagreement is not randomly distributed, but instead concentrated in protein classes with limited annotation, complex structure-function relationships, or atypical evolutionary trajectories.

### Residue-level structural context shapes concordance among VEPs

Building on the protein-class trends described above, we next asked whether residue-level structural context could explain where and why VEPs disagree, and whether one or more structural signals could be used to prioritise genetic targets for experimental investigation. Using sequence-matched AlphaFold2 models from the AlphaFold Protein Structure Database (Fig. 4A), we annotated every residue in each structure with a uniform set of descriptors: model confidence (pLDDT) as a proxy for intrinsic-disorder propensity (***Akdel et al., 2022***), solvent exposure (rASA (***Rost and Sander, 1994***) and its complementary weighted-contact number, WCN), and secondary-structure assignments (Fig. 4B). Because each protein had already been assigned to a low-, intermediate-, or high-concordance tier based on the median Spearman correlation coefficient of polarity-aligned VEP scores across shared SNVs (Fig. 3A), we propagated these protein-level labels to their residues and summarised each structural descriptor by its median value per protein for downstream comparisons (Fig. 4B).

**Figure 4.**
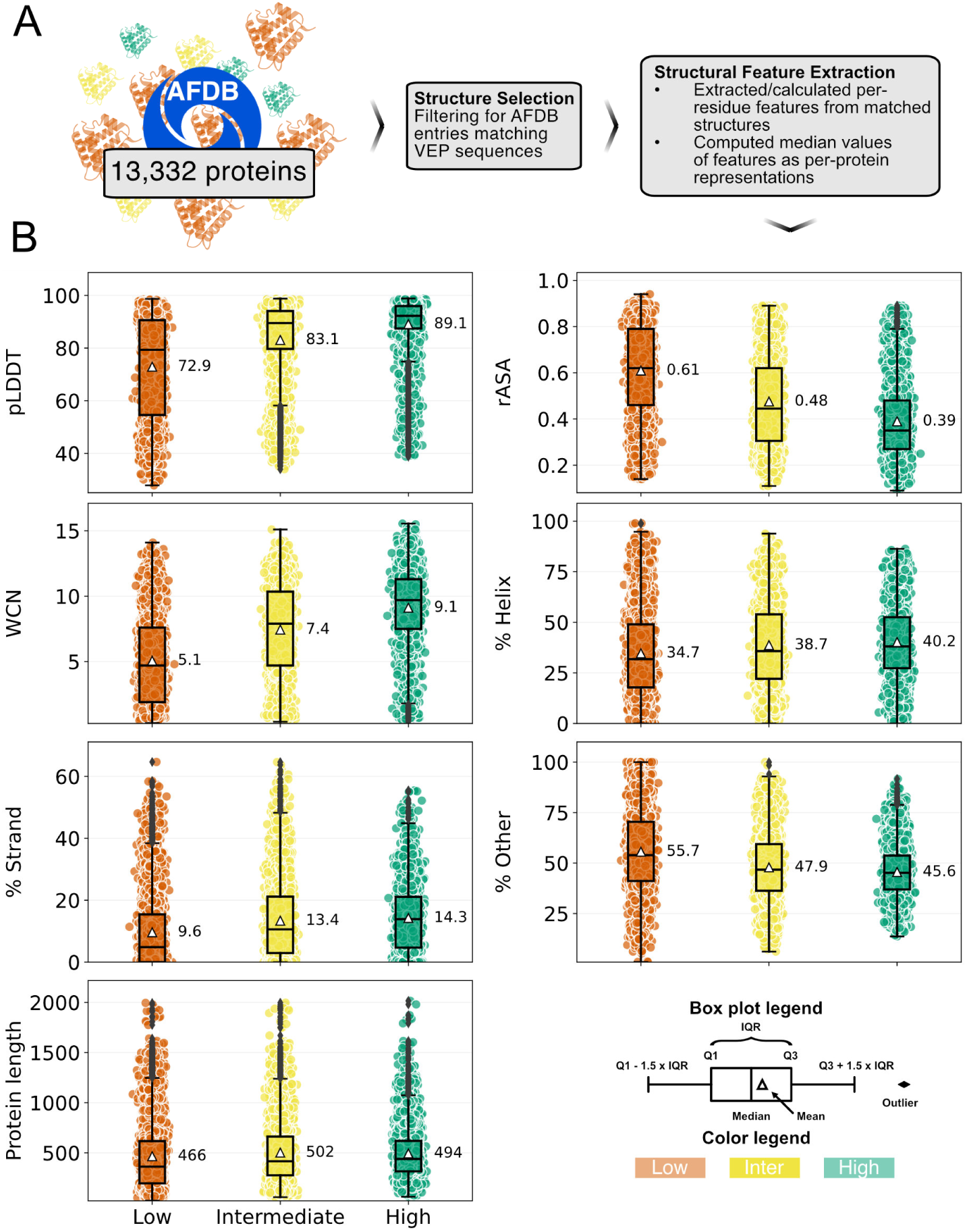
Protein-level structural features derived from sequence-matched AlphaFold2 models. (A) Structure selection and matching: AlphaFold Protein Structure Database entries whose sequences exactly match those used by the predictors were retrieved to avoid numbering offsets and coverage artefacts. (B) Feature extraction and summarization: per-residue metrics—model confidence (pLDDT), relative solvent accessibility (rASA) with its complementary weighted contact number (WCN), and secondary-structure assignments—were computed for each model and summarised by their median values to yield a single representative score per protein for downstream analyses.

A central outcome of this analysis is that the main structural features associated with inter-VEP concordance point to the same underlying axis, which runs from confidently modelled, well-packed cores to flexibly modelled, surface-exposed, and often intrinsically disordered segments, as shown in prior work linking low pLDDT to disorder and increased exposure (***Akdel et al., 2022***; ***Fawzy and Marsh, 2025***; ***Tunyasuvunakool et al., 2021***). This matters for a practical reason. If VEP disagreement is concentrated in a specific residue context, then the same context can be used to flag variants where experimental follow-up is most likely to resolve uncertainty. Consistent with this, proteins in the low-agreement tier show markedly lower median pLDDT, often below the conventional disorder threshold of 70, together with higher exposure and less regular secondary structure (Fig. 4B). In contrast, high concordance concentrates in proteins dominated by confidently modelled proteins with high median pLDDT, buried residues (low rASA and high WCN), and higher fractions of regular secondary structure (Fig. 4B). The simplest mechanistic interpretation is that disordered and flexible regions lack stable tertiary constraints, which weakens both stability-related evidence and alignment-based conservation signals, as shown in studies of disorder and variant-effect prediction (***Cagiada et al., 2025***; ***Fawzy and Marsh, 2025***; ***Uversky et al., 2008***). These segments are also under-represented in crystallographic training data and are often shaped by context-specific constraints such as transient interactions or short regulatory motifs, which makes predictors trained primarily on ordered domains less reliable in this regime, as shown in analyses of training bias and context dependence (***Fawzy and Marsh, 2025***; ***Livesey and Marsh, 2020***). When the usual cues are weak or inconsistent, different VEP families fall back on different assumptions. Evolutionary-depth methods such as GEMME and ESCOTT can produce attenuated or noisy estimates when alignments are shallow or signals are patchy, whereas language-model approaches can still assign high deleteriousness when local sequence context resembles constrained patterns (***Brandes et al., 2023***; ***Meier et al., 2021***). VEPs that are trained on clinical data or tuned on population-level data may further up-weight exposed substitutions when they overlap with known disease-relevant motifs or hotspots, while stability-focused VEPs often treat many surface substitutions as benign (***Fawzy and Marsh, 2024***; ***Livesey and Marsh, 2025***; ***Findlay et al., 2018***). Seen this way, what can look like separate contributions of disorder, surface exposure, and reduced secondary structure is perhaps better treated as one residue context where structural and evolutionary information is less decisive, which in turn magnifies differences among predictors, as argued in prior work on inter-predictor discordance (***Fawzy and Marsh, 2025***; ***Brandes et al., 2023***). This interpretation aligns with reports that low-concordance predictor clusters are enriched in IDR-heavy settings where evolutionary constraints are weaker, and that exposure-related metrics and coil enrichment track reduced agreement (***Fawzy and Marsh, 2024***, ***2025***; ***Brandes et al., 2023***). Protein length varies only modestly across concordance tiers, with median lengths 466, 502, and 494 residues for low-, intermediate-, and high-concordance proteins, respectively (Fig. 4B), indicating that overall size is secondary to whether proteins are dominated by ordered cores or flexible, exposed segments, as shown in the tier-wise comparisons (***Fawzy and Marsh, 2025***).

Taken together, these results support a simple unifying principle. Residue-level structural context can be used both to explain where VEPs disagree and to prioritise experimental targets, because concordance is highest when evolutionary, structural, and biochemical constraints align in ordered, buried, confidently modelled regions, and lowest when those cues become ambiguous in exposed, flexibly modelled, coil-rich segments, as shown in prior disorder-focused analyses (***Akdel et al., 2022***; ***Fawzy and Marsh, 2025***). This framing also reconciles residue-level mechanisms back to the protein-class trends. Proteins enriched in flexible linkers and disordered loops shift toward low concordance, whereas proteins dominated by rigid secondary structure and packed cores tend to populate the high-agreement regime. Methodologically, it suggests that reducing disagreement will require better representation and explicit modelling of the flexible-surface and IDR regime, including broader training coverage, motif-aware features for surface interaction sites, and hybrid models that combine language-model embeddings with physics- or motif-informed priors, as shown in recent proposals and evaluations (***Tesei et al., 2024***; ***Voutsinos et al., 2025***; ***Larsen et al., 2025***). For IDRs in particular, progress may depend on combining ensemble descriptions (***Tesei et al., 2024***), degron maps (***Voutsinos et al., 2025***; ***Larsen et al., 2025***), IDR-oriented language models (***Mollaei et al., 2024***), and alignment-free approaches that remain informative when evolutionary context is sparse (***Pritišanac et al., 2024***; ***Ruff et al., 2026***).

### Population-genetic constraint is associated with inter-VEP concordance

Beyond structural descriptors, we asked whether gene-level signatures of selective constraints could further highlight why some proteins show markedly divergent VEP scores whereas others do not. To this end, we mapped the proteins in our concordance panel to gnomAD-derived constraint metrics. The probability of loss-of-function intolerance (pLI), its complementary probabilities of recessive tolerance (pRec) and neutrality (pNull), the missense Z-score, and lastly the heterozygous selection coefficient *S*_het (***Zeng et al., 2024***) (Fig. 5). As with the structural analysis, these values were compared across the low-, intermediate-, and high-agreement tiers that had been established from VEP correlations.

**Figure 5.**
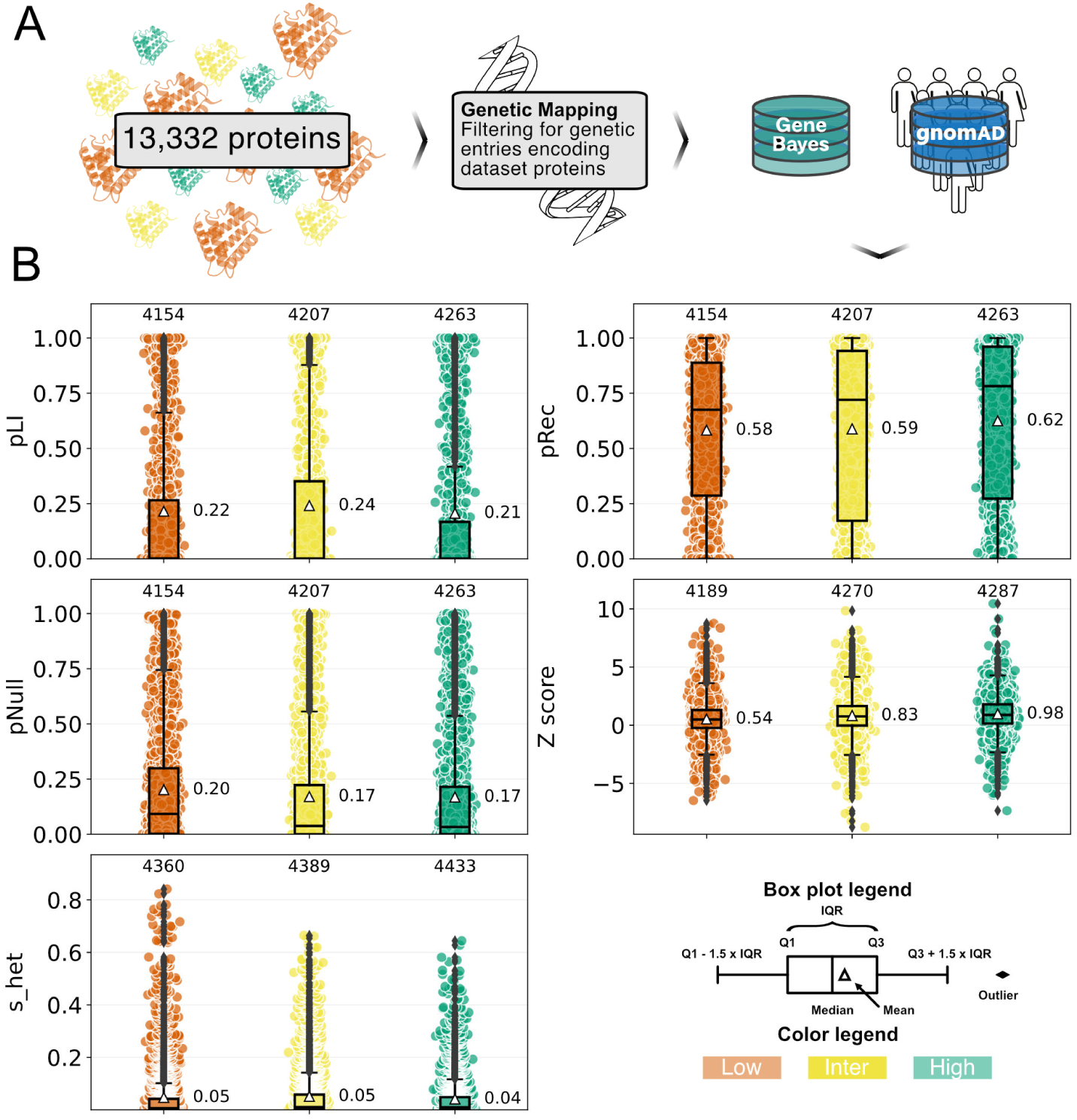
Population-genetic and gene-level constraints associated with inter-VEP concordance. (A) Population-genetic constraints extracted from gnomAD (***Karczewski et al., 2020***) and GeneBayes (***Zeng et al., 2024***) databases and filtered for matching inter-VEP concordance protein targets. (B) Gene-level metrics from gnomAD—missense Z-score, pLI, pRec, pNull, and the heterozygous selection coefficient (*S*_het_)—were compared across the low-, intermediate-, and high-agreement tiers. Distributions show that missense Z and pRec increase from low to high agreement, consistent with stronger depletion of missense variants in high-concordance genes, whereas pLI and *S*_het_ display weak or inconsistent separation; pNull mirrors the inverse of pRec. Because the constraint databases are incomplete, matched constraint data were available for different numbers of proteins depending on the model, as annotated above each category in the subplot.

Overall, we find substantial overlap across classes, showing that these metrics are not tightly coupled to the inter-VEP consistency classes. Beyond that, two broad patterns emerged. First, metrics that capture coding constraint at the amino acid level were the most discriminative. The missense Z-score, which quantifies depletion of missense variants relative to expectation, increased monotonically from low-through intermediate- to high-agreement proteins (0.54, 0.83, and 0.98, respectively). This finding is consistent with prior observations that VEP concordance is higher in genes and domains under stronger purifying selection, where evolutionary signals are more consistent across prediction methods (***Fawzy and Marsh, 2024***; ***Livesey and Marsh, 2020***, ***2025***). High missense Z-scores often correspond to ordered domains with well-defined tertiary structure and high AlphaFold2 per-residue confidence, in which residue-level features such as solvent accessibility and catalytic site geometry are reliable predictors of functional impact (***Varadi et al., 2022***; ***Akdel et al., 2022***; ***Fawzy and Marsh, 2024***). Likewise, pRec—the probability that a gene tolerates loss-of-function only in the homozygous state—was highest among high-agreement proteins (0.62 versus, 0.58, and 0.59 in low- and intermediate-agreement groups, respectively). This aligns with the enrichment of high-concordance targets among recessive disease genes, which frequently harbour highly conserved structural or catalytic domains (***Fawzy and Marsh, 2024***; ***Livesey and Marsh, 2025***). In such contexts, mutational constraints are clearer and more uniform, making them easier for predictors to capture (***Fawzy and Marsh, 2025***). Second, genome-wide measures of general haploinsufficiency or heterozygous intolerance showed comparatively weak or inconsistent relationships with VEP agreement. Mean pLI values were indistinguishable across tiers, and the heterozygous selection coefficient *S*_het differed by only one hundredth between groups. This muted separation mirrors earlier findings that global dosage sensitivity at the gene level is less predictive of concordance than residue-level constraints (***Fawzy and Marsh, 2024***; ***Livesey and Marsh, 2025***). Together, these trends imply that predictors diverge more on the qualitative nature of amino acid substitutions within a gene—shaped by local structural order, motif content, and evolutionary conservation—than on the overall gene-level sensitivity to variation (***Fawzy and Marsh, 2024***, ***2025***; ***Livesey and Marsh, 2025***).

Taken together with the structural context analysis, the genetic constraint data reinforce a consistent theme that VEP concordance is highest when multiple, orthogonal signals of variant intolerance converge. These include strong evolutionary conservation within ordered domains—often accompanied by high AlphaFold2 per-residue confidence—biophysical indispensability at catalytic sites, binding pockets, or other structurally integral residues, and population-level depletion of missense variants. Notably, gene-level summaries such as pRec, pLI, and *S*_het_ are inferred from observed-versus-expected patterns across the whole gene and as such do not label individual variants but reflect a gene’s overall tendency toward constraint (e.g., recessive tolerance for pRec). However, when these gene-level tendencies align with local, residue-level evidence, they define a coherent constraint landscape in which predictors with different architectures tend to agree. Conversely, concordance is lowest where one or more signals is weak, heterogeneous, or absent— such as in IDR-rich proteins, non-canonical segments, or genes with mixed domain architectures and variable tolerance profiles.

### Predictor consensus does not correlate with experimental MAVE measurements

Having established protein class-specific patterns alongside structural and population-genetic effects, we next asked whether cross-model consensus aligns with independent experimental benchmarks. We focused on MAVEs, which provide dense, quantitative maps of functional consequences for thousands of single amino acid substitutions in a single experiment and thus serve as a benchmark for prediction models. We note here that the ten VEPs analysed here were chosen because they show comparatively good agreement with experimental datasets overall. If inter-VEP agreement reflects the same underlying biological signal that MAVEs capture, then proteins with high cross-model agreement should, on average, show stronger correlations between VEP scores and MAVE readouts than proteins with low agreement. Conversely, a weak or absent relationship would imply that the factors driving computational concordance are not, in general, the factors that determine performance on these experimental assays, revealing potential ‘blind spots’ in current predictive frameworks and underscoring the need for targeted experimental follow-up.

To test this hypothesis we used a previously curated set of 36 high-quality MAVE datasets (***Livesey and Marsh, 2025***), selecting one representative MAVE readout per protein (see Methods). After harmonising residue numbering schemes and restricting to substitutions represented in both the MAVE and the ten-predictor subset, we derived two orthogonal per-protein summaries: (i) inter-VEP concordance, defined as the median absolute Spearman correlation across all VEPs after polarity alignment of predictor scores (Fig. 3A) and (ii) prediction accuracy, defined as the median absolute Spearman correlations between the MAVE readout and each VEP. All correlations for a given protein were evaluated on the same MAVE-anchored shared variant set—i.e., variants with non-missing scores for the selected representative MAVE readout and all included VEPs—ensuring comparability between inter-VEP and VEP-MAVE metrics. Absolute Spearman correlations were used to capture rank agreement independent of score polarity or calibration, and medians were used to reduce sensitivity to outliers or individual predictor behaviour. We note that seven proteins were excluded after representative MAVE selection because at least one of the ten VEPs lacked predictions for the assayed variants, leaving *n* = 29 proteins for the analysis. We summarised these results in a scatter plot of inter-VEP concordance versus prediction accuracy to assess whether inter-VEP predictor agreement co-varies with agreement to experimental measurements (Fig. 6). For transparency, we provide per-protein summaries of VEP-MAVE correlations across all predictors in Supplementary Fig. S2; these panels support the condensed analysis. Our results show no clear relationship between predictor consensus and agreement with experiment. Notably, we found that 69% of proteins in the MAVE benchmark (20 of 29) fall within the high inter-VEP concordance category, even though this category comprises only one third of proteins in the concordance distribution. This indicates that VEPs may have biased accuracies for proteins with MAVEs, or that proteins selected for MAVEs constitute a biased subset. We note here that this observation may also be biased by our choice of VEPs that, on average, show high agreement with MAVEs. Nevertheless, we find that proteins with very similar inter-VEP concordance can occupy widely different positions on the experimental predictability axis.

**Figure 6.**
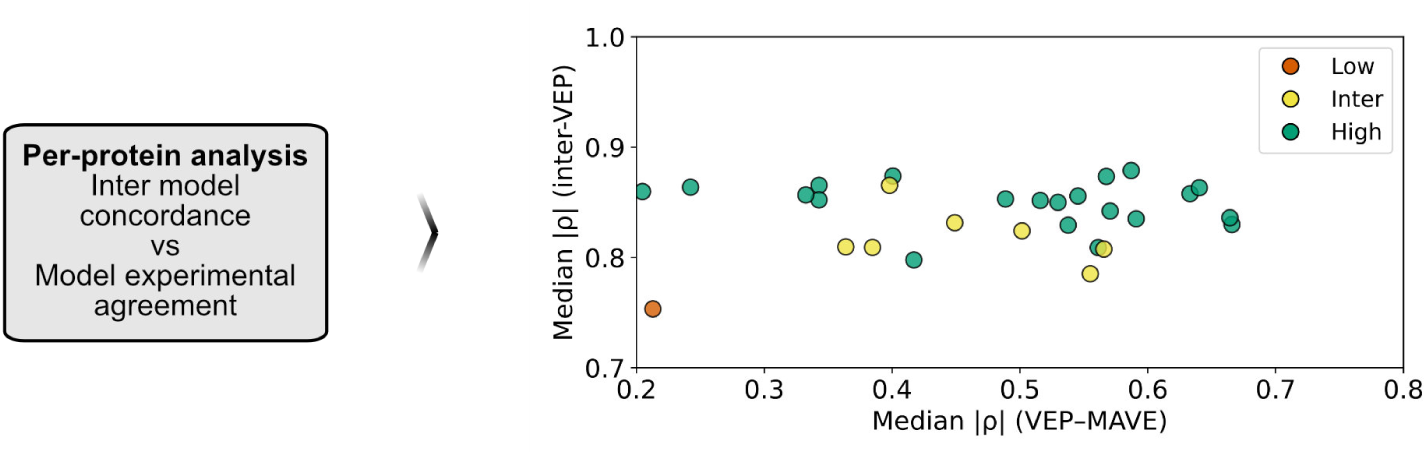
Computational-experimental comparison across MAVE datasets. Each point represents one of 29 proteins characterised by a MAVE. For each protein, correlations were computed on a shared set of variants containing only SNVs with available scores for both the selected VEPs and the chosen MAVE dataset. The y-axis shows inter-VEP concordance—defined as the median pairwise absolute Spearman correlation among the ten selected VEPs after polarity alignment of predictor scores computed on this shared variant set—while the x-axis shows prediction accuracy, defined as the median absolute Spearman correlation between the MAVE results and each VEP, also computed on the same shared variant set. Points are coloured according to previously defined agreement categories (low, intermediate, high), which were determined by dividing the distribution of median inter-VEP correlations across proteins into three equally sized groups. The results show that strong predictor consensus does not necessarily imply alignment with empirical fitness maps.

This decoupling has a direct interpretation for this benchmark panel. The scatter plot is dominated by substantial variation in prediction accuracy at broadly similar (often high) levels of inter-VEP concordance, implying that models frequently ‘succeed together’ or ‘fail together’ on a given protein. This yields two empirically distinct regimes with different practical meanings. In high-consensus/high-validity proteins (e.g., PRKN, inter-VEP |*ρ*| = 0.857, VEP-MAVE |*ρ*| = 0.63), multiple predictors not only agree with each other but also recover the experimental ranking of variant effects, suggesting that the information shared across models is assay-relevant for that target. In contrast, high-consensus/low-validity proteins (e.g., HMGCR, inter-VEP |*ρ*| = 0.865, VEP-MAVE |*ρ*| = 0.37) represent cases where predictors converge on a shared prioritisation yet collectively fail to match the experimental readout. These proteins are particularly informative because the failure mode is shared across models and therefore cannot be diagnosed by cross-model agreement alone.

The lack of correlation indicates that predictor consensus does not guarantee experimental validity and, by extension, that current VEPs and MAVEs are probing only partially overlapping facets of protein biology. A parsimonious explanation is that inter-VEP concordance is often driven by partially overlapping information sources and inductive biases common to many predictors— most prominently evolutionary conservation and physicochemical substitution propensities, and in some cases population-genetic priors—which can induce strong model-to-model agreement regardless of whether those signals are sufficient to capture the assay endpoint (***Livesey and Marsh, 2025***; ***Fawzy and Marsh, 2024***). These conservation and biophysics-based priors can be highly predictive in well-ordered domains with strong evolutionary constraints but tend to offer less sensitivity in regions where functional impact is mediated by context-specific mechanisms (***Fawzy and Marsh, 2025***, ***2024***). In parallel, the MAVE-derived datasets in this panel do not measure a single unified phenotype. Each assay reports variant effects in a specific experimental context (e.g., growth/fitness proxies, interaction disruption, or abundance-related readouts), and different contexts can weight different mechanistic determinants of variant impact (***Findlay et al., 2018***; ***Starita et al., 2017***). Consistent with prior work, predictor performance can therefore vary by assay modality and target even when inter-model concordance remains high (***Fawzy and Marsh, 2025***). For instance, MAVEs that probe protein abundance are more strongly correlated with VEPs that probe structural stability whereas functional assays are better predicted by VEPs that use sequence conservation (***Jepsen et al., 2020***; ***Cagiada et al., 2021***). However, we emphasise that Fig. 6 demonstrates the decoupling itself; attributing any individual low-validity case to a particular mechanism requires assay- and target-specific follow-up.

These results suggest two practical implications for interpreting VEP consensus and for prioritising experiments. First, high concordance among VEPs should not be interpreted as a ‘solved’ target; in this panel, consensus is common even where agreement with experiment is modest, indicating that it can arise from shared priors rather than mechanistic correctness (***Livesey and Marsh, 2025***; ***Fawzy and Marsh, 2024***). Empirical validation through appropriately chosen functional assays therefore remains essential, including for proteins with strong predictor consensus (***Starita et al., 2017***). Second, experimental resources are likely to yield the greatest informational return when allocated to targets characterised by elevated model uncertainty or by features that may limit the reliability of current predictors. Predictor disagreement can be identified prospectively and highlights where model assumptions diverge, marking regions where computational evidence varies. In addition, our analysis suggests that certain protein classes and mechanistic contexts are more prone to systematic prediction errors, even when inter-VEP concordance is high. Although true blind spots can only be confirmed with experimental data, these patterns provide guidance on which types of proteins may benefit most from targeted functional assays. Prioritising such targets is expected to reduce variants of uncertain significance and improve clinical interpretability by resolving regions where current computational evidence is least informative or systematically biased (***Fawzy and Marsh, 2024***; ***Findlay et al., 2018***; ***Dace et al., 2025***). Guided by these principles, we next outline how our observations may inform the selection of proteins for experimental characterization.

### Using predictor discordance to prioritise targets for experimental investigation

The finding that inter-VEP concordance does not correlate with agreement to MAVE measurements has implications for how experimental resources should be allocated. High predictor consensus cannot be assumed to reflect biological accuracy; it may instead reflect shared methodological priors (e.g., reliance on evolutionary conservation metrics, allele-frequency constraints, or physicochemical substitution heuristics) rather than a faithful capture of the molecular mechanisms underlying variant effects (***Fawzy and Marsh, 2024***; ***Livesey and Marsh, 2025***). In contrast, divergence among VEPs can be informative because it highlights proteins where predictors disagree most, which may reflect mechanisms the models do not capture well and therefore makes these proteins strong candidates for experimental investigation.

This observation motivates a prioritization funnel that starts broadly and narrows to a concise shortlist of candidates for future experimental follow-up (Fig. 7). Evidence that the gene is important in human cells—such as essentiality or strong fitness effects—could further support prioritization by increasing the likelihood that variant effects produce measurable cellular consequences. In practice, we begin by ranking proteins using the inter-VEP concordance metric introduced earlier and assigning them to low-, intermediate-, and high-agreement tiers using fixed quantile thresholds. The low-agreement tier forms the initial pool because it is enriched for targets that expose limitations in current prediction frameworks and, as shown by our benchmarking with data generated by MAVEs, is not simply reproducing patterns already captured by proteins with existing MAVE data. However, our residue-level structural analysis indicates that much of the low-concordance tier is explained by one dominant regime—proteins enriched in disordered or flexibly modelled, surface-exposed, coil/loop segments (low pLDDT, high rASA/low WCN, and increased coil content). Because this regime is a well-documented difficulty for VEPs and is also frequently recalcitrant to conventional high-throughput assays—whose readouts (e.g., growth rescue or simple reporter activity) often miss the subtle, context-dependent effects that dominate disordered, surface-exposed, or coil-rich regions (***Fawzy and Marsh, 2025***; ***Starita et al., 2017***)—we refine the low-agreement pool by separating proteins whose low concordance is likely driven by extensive flexible or disordered surface composition from those that remain low-concordance despite being confidently modelled and structurally ordered. The latter group is particularly attractive for experimental investigation because it is more tractable and disagreement is less likely to be explained by disorder-related ambiguity alone, making it more likely to point to specific missing mechanisms.

**Figure 7.**
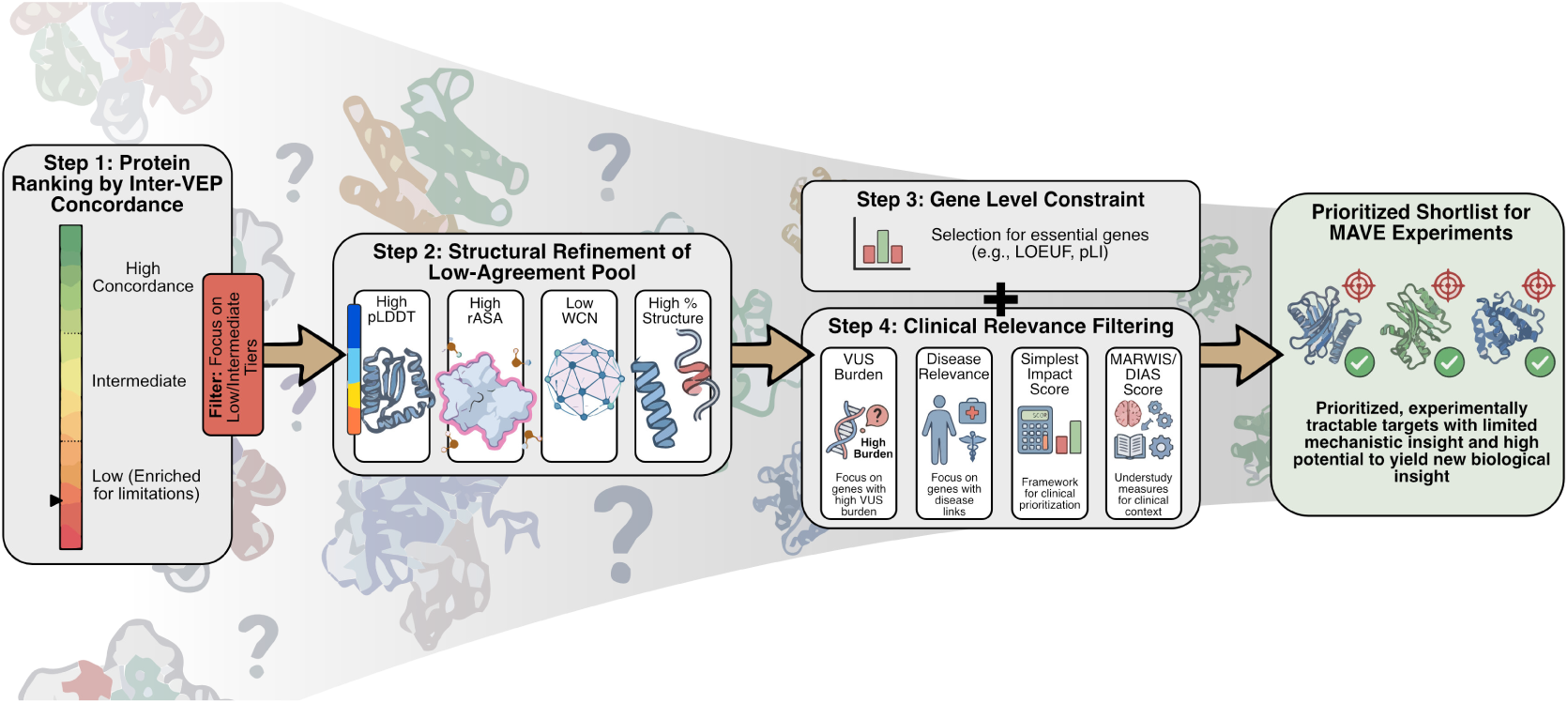
Prioritization funnel for selecting proteins for empirical investigation. Proteins are first ranked by inter-VEP concordance and low-agreement targets are retained as an initial pool enriched for limitations in current prediction frameworks. This pool is then structurally refined using AlphaFold2-derived features (pLDDT, rASA, WCN, and secondary-structure composition) to distinguish proteins whose discordance is driven by extensive flexible or disordered surface regions from those that remain low-concordance despite confident structural ordering. Gene-level constraint metrics (e.g., LOEUF, pLI) provide a second prioritization axis, followed by clinical relevance filters incorporating VUS burden, disease association, Simplest Impact Score, and MARWIS. Together, these sequential filters yield a shortlist of experimentally tractable, clinically meaningful candidates for future MAVE studies.

Operationally, we propose using the AlphaFold2-derived summaries already described above to refine the initial low-concordance protein set. Gene-level constraint metrics can then be added as a secondary axis. Accordingly, low-concordance, structurally ordered targets in more constrained genes represent plausible high-value candidates, whereas low-concordance targets in weakly constrained genes may offer lower yield for generalizable mechanism or clinically relevant reclassification (***Fawzy and Marsh, 2024***, ***2025***; ***Livesey and Marsh, 2025***). A final prioritization layer incorporates clinical relevance by focusing on genes with substantial VUS burdens in ClinVar and clear disease links. Existing frameworks support this step, including the Simplest Impact Score (ranking by unique missense VUS counts after excluding already classified variants), MARWIS (up-weighting “movable” VUS and variants that recur across unrelated patients), and DAIS (the difficulty-adjusted impact score which further accounts for experimental tractability when ranking targets) (***Kuang et al., 2021***), as well as knowledge-based measures of how under-studied a gene is (***Kuang et al., 2021***; Chen et al., 2022). Applying these filters after the structural and constraint triage yields a concise, assay-ready shortlist that is both experimentally tractable and directly aligned with the goal of reducing VUS uncertainty in clinically important genes.

As a concrete illustration, we begin with proteins that exhibit low inter-VEP concordance and then prioritise those that are structurally well resolved and therefore likely more straightforward to assay and interpret. Because our earlier analyses show that low pLDDT, high rASA/low WCN, and elevated coil fraction largely track the same underlying regime of disorder and flexible architecture, we use a high median AlphaFold2 pLDDT score as a filter for structural tractability. A high pLDDT threshold (e.g., >90) increases confidence that the protein adopts a well-defined structure, and since MAVE studies of IDRs often show weakly position-specific effects that mainly reflect overall composition or amino-acid class (***Shepherdson et al., 2024***; ***Staller et al., 2018***), focusing on high-pLDDT targets should yield fitness-assay results that are more site-specific and easier to interpret in structural terms. Under this criterion, the transporter-like protein OSCP1 (*OSCP1*, UniProt Q8WVF1) emerges as a compelling candidate. OSCP1 shows low VEP agreement (inter-VEP |*ρ*| = 0.29), yet its median pLDDT score is high (≈ 92), indicating that it could be experimentally tractable. Moreover, although OSCP1 has been reported to function as a putative organic solute carrier involved in placental drug clearance (***Kobayashi et al., 2005***), its broader physiological role and mechanistic context remain incompletely defined, and as of 2025-01-02 ClinVar records 35 variants of uncertain significance and only a single benign interpretation, providing a rationale for targeted functional mapping. In this way, disagreement among computational predictors becomes a practical signal for selecting experiments that can improve mechanistic understanding while supporting variant interpretation and reducing the VUS burden.

## Conclusions

Large-scale comparison of VEPs across the human proteome reveals a structured, heterogeneous landscape of agreement. Clustering of 71 tools, and a focused examination of ten methodologically diverse predictors, shows that concordance is shaped by modelling lineage and shared input features—not scattered at random. At the protein-class level, low agreement is enriched in categories where the signals most VEPs rely on are less informative or less transferable, including proteins of viral/transposable-element origin and immune/defense proteins (often shaped by rapid diversification), as well as structural and storage proteins whose functional constraints can be weakly captured by standard conservation- and monomer-centric feature sets. At the residue level, reduced AlphaFold2 confidence (lower median pLDDT), higher solvent exposure, and coil/loop-rich secondary structure align with poorer concordance, whereas buried, confidently modelled cores yield much higher agreement. Population-genetic patterns support this observation but correlate more weakly with concordance. Genes depleted for missense variants show slightly elevated concordance, while genome-wide proxies for haploinsufficiency provide little discrimination. Importantly, these gene-level summaries reflect a gene’s overall tendency toward constraint rather than labelling individual variants.

We also find that inter-VEP concordance bears no consistent relationship to empirical measurements from MAVEs. Across the curated MAVE panel, strong predictor consensus does not imply stronger alignment with experimental readouts; conversely, proteins marked by disagreement are just as likely to expose functionally consequential discrepancies. This decoupling cautions against equating consensus with correctness. Agreement can reflect shared priors and shared biases as easily as biological truth, while disagreement can flag contexts where current feature sets and training regimes diverge in their assumptions. These results therefore shift the logic of experimental triage. We suggest to prioritise targets where predictors disagree—especially cases that remain low-concordance despite having structurally well-resolved AlphaFold2 models, where disagreement is less likely to be explained by pervasive disorder alone, and where follow-up is more interpretable. From this premise, we propose a simple selection funnel: (i) rank proteins by low inter-VEP concordance; (ii) enrich for structurally tractable targets using protein-level AlphaFold2 features; and (iii) optionally refine using gene-level constraint and clinical-variant landscape measures. Nonetheless, unanticipated practical constraints may still limit experimental investigation for some targets. Prioritised targets should ultimately be assessed for tractability, including whether a practicable quantitative readout exists (e.g., activity, binding/interaction, localization, transport, abundance/fitness proxy) in a suitable system. Accordingly, experimental feasibility is a downstream consideration rather than an output of the present study.

One further implication concerns the choice of experimental readout. Our analyses treat MAVEs as a unified source of empirical information, but different assay modalities probe distinct molecular consequences of variation, including effects on protein stability, abundance, interaction networks, localisation, or catalytic activity. Disagreement among predictors may often arise because different models implicitly prioritise different mechanistic determinants of variant impact. In such cases, experimental assays that probe multiple complementary phenotypes, or that are selected to test specific mechanistic hypotheses suggested by predictor divergence, may yield the greatest informational return. While the present study does not attempt to prescribe specific assay types for individual targets, aligning experimental design with the likely sources of predictor disagreement represents a promising strategy for maximising the value of future MAVE datasets.

These considerations should be interpreted in the context of several important limitations. Concordance rankings depend on the chosen predictor panel and the harmonised variant set; alternative datasets and coverage profiles may shift local rankings even if broad trends persist. Thus, some of our conclusions may be biased by selecting VEPs that show overall high agreement with existing MAVEs. Protein-level structural summaries can mask domain-level heterogeneity and interface-specific effects, and MAVE readouts vary by system and phenotype, so correlations between predictor scores and experimental effects are not directly comparable across datasets. Finally, while our analyses span thousands of proteins, they cover only a subset of human proteins and variants that are jointly scored in the harmonised datasets used here, leaving additional targets outside the present comparison. Within these bounds, inter-VEP disagreement nonetheless emerges as a practical signal for prioritizing experimental resources and a generalizable route for converting computational uncertainty into experimental action—effectively expanding and diversifying functional assay coverage, strengthening benchmarking, and guiding the next generation of VEP models as datasets grow in breadth and mechanistic resolution.

## Methods

### Construction of the genome-wide VEP concordance matrix

To quantify the similarity of variant effect predictors, we compared all pairs of the 71 VEPs (Table 1). Raw scores were downloaded from dbNSFP (***Liu et al., 2020***), project-specific APIs, or generated locally from the released source code using default settings. Predictor scores were polarity-aligned so that larger values consistently reflected greater predicted deleteriousness, following the authors’ documentation or, when ambiguous, the polarity that maximised agreement with benchmark multiplexed assays (see Supplementary Methods). A fully harmonised ‘all-shared’ matrix was obtained by retaining only those single-nucleotide variants for which every predictor returned a score, yielding 6,805,018 missense SNVs that map to 1,242,248 residue positions in 3,846 unique proteins. We then calculated the Spearman’s rank correlation coefficient, *ρ_ij_*, for each pair of predictors across this variant set using the tie-aware `scipy.stats.spearmanr` function (v1.12.0), producing a signed 71 × 71 correlation matrix. To compare predictors at the level of their global concordance patterns, the correlation matrix was interpreted as a profile representation in which each VEP is described by the vector of its correlations with all other predictors. In this formulation, similarity between predictors reflects the degree to which they agree with the same subset of methods across the broader ensemble, rather than the magnitude of their direct pairwise correlations alone. Distance between predictors was therefore computed using correlation distance applied to these correlation profiles, such that two VEPs are considered similar when their patterns of agreement with the remaining predictors follow a comparable ranking structure.

We performed an agglomerative hierarchical clustering using average linkage, implemented via `seaborn.clustermap` calling `scipy.spatial.distance.pdist(metric=`“`correlation`”) and `scipy.cluster.hierarchy.linkage(method=`“`average`”). Average linkage iteratively merges clusters based on the mean pairwise distance between their members, producing a hierarchy that reflects overall similarity in correlation profiles while remaining robust to local fluctuations in individual pairwise correlations. Using correlation distance in this context emphasises shared agreement structure across the predictor ensemble and is insensitive to uniform shifts in correlation magnitude, allowing predictors to cluster together when they exhibit similar concordance patterns even if the strength of individual pairwise correlations differs. The resulting dendrogram and clustered heatmap, rendered using a diverging colour scale spanning the full correlation range (−1 to 1), are shown in Fig. 2 and provide the basis for the global concordance analysis discussed in the main text.

To verify that restricting analysis to universally scored variants did not distort the concordance landscape, we repeated the calculation on a pair-specific basis, recomputing *ρ_ij_* for each VEP pair using the maximally overlapping set of variants jointly scored by that pair (including multi-nucleotide substitutions where available). The resulting pair-specific correlation matrix showed the same qualitative concordance structure as the all-shared matrix (Supplementary Fig. S1), confirming that clustering patterns are robust to the variant inclusion strategy. For all subsequent analyses—including the protein-level concordance metrics described above—we retained the all-shared correlation matrix to eliminate coverage-related confounders and maximise interpretability.

### Protein-level concordance metrics

Inter-VEP agreement was quantified for every canonical (UniProt) protein represented in the ten-predictor, all-shared SNV missense dataset, which comprised 37,213,055 scored variants spanning 6,333,694 distinct residue positions in 13,332 proteins. Because we filtered for SNVs where all ten selected predictors—AlphaMissense, CPT-1, GEMME, ESCOTT, ESM-1v, popEVE, VARITY_R, MetaRNN, SNPred, and gMVP—returned a value for every variant in this set, pairwise comparisons could be performed without imputation or additional coverage filtering. For a given protein *p* we assembled an *n_p_* × 10 matrix of predictor scores, where *n_p_* is the number of shared single-nucleotide variants mapped to that protein; no minimum *n_p_* threshold was imposed. Spearman’s rank correlation coefficient, calculated with the tie-aware `scipy.stats.spearmanr` function (v1.12.0), was chosen because it captures monotonic relationships and is robust to the heterogeneous score distributions produced by different algorithms. For each of the 45 predictor pairs (*i, j*) we computed the correlation 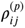 across the *n_p_* variants after polarity alignment of predictor scores and summarised the 45 resulting values by their median to obtain a single concordance score *C_p_* for that protein. The distribution of *C_p_* across all 13,332 proteins was broad and continuous. For downstream comparisons we discretised this distribution into three equally sized concordance tiers (low, intermediate, and high) using the empirical 0.33 and 0.67 quantiles of *C_p_* as cut-offs (*C*_0.33_ = 0.7685 and *C*_0.67_ = 0.8242). Proteins with *C_p_* < *C*_0.33_ (*n* = 4, 444) were classified as low agreement, those with *C*_0.33_ ≤ *C_p_* ≤ *C*_0.67_ (*n* = 4, 444) as intermediate agreement, and those with *C_p_ > C*_0 67_ (*n* = 4, 444) as high agreement. These tier assignments were used as categorical predictors in enrichment analyses, while the continuous *C_p_* values were retained for analyses treating concordance as a continuous variable.

### Protein functional class annotation

To determine whether predictor disagreement clusters within particular functional categories, we first annotated every protein in the concordance panel with a top-level protein-class label from the PANTHER database (release 19.0). The PANTHER hierarchy comprises a manually curated tree in which the 24 root classes—such as *metabolite-interconversion enzyme*, *cytoskeletal protein* and *viral or transposable element protein*—sit above other nested child terms. Because the downloadable flat files do not expose the full ancestry of each leaf node, we reconstructed the hierarchy by programmatically scraping the category pages at https://www.pantherdb.org/panther/category.do? categoryAcc=PC00000, iterating through every child link, and recording all parent-child relationships. The resulting adjacency list was converted into a directed acyclic graph, and a reverse mapping was derived so that every leaf term points unambiguously to exactly one of the 24 top-level classes.

UniProt accessions were then linked to leaf-level protein-class identifiers using the directed acyclic graph we constructed from the v.19.0 release of the PANTHER DB. Each leaf term was propagated upward through the reconstructed graph, yielding a single root-class label for each protein; duplicate leaf assignments that resolved to the same root were collapsed, whereas genuine conflicts between different roots resulted in multiple class assignments. This procedure produced a complete contingency table of the 13,332 proteins versus the 24 PANTHER root classes plus the *unknown* bin. Finally, these class labels were intersected with the previously defined low-, intermediate-, and high-concordance tiers; within each root class we counted proteins in the three tiers and visualised the distribution (Fig. 3B).

### Structural feature extraction from AlphaFold models

For every protein in the concordance panel we obtained a corresponding structural model from the AlphaFold Protein Structure Database (AFDB, version 4, accessed April 2025) (***Varadi et al., 2022***). Retrieval began with the ‘one-sequence-per-gene’ reference proteome for *Homo sapiens* (UniProt proteome UP000005640, release 2021_04), the same sequence collection used to generate the AFDB human set. UniProt accessions in the concordance panel were intersected with this list; entries that failed any of the AFDB preprocessing criteria-sequence length shorter than 16 residues, presence of non-standard amino acids, or absent from the AFDB reference list were discarded. AFDB predictions for proteins longer than 2,700 residues are supplied as partially overlapping 1,400-residue segments; for such proteins we extracted per-residue pLDDT and solvent-exposure values from each segment and averaged the 1,200-residue overlaps position-wise to obtain a single continuous profile.

Downloaded AlphaFold2 coordinate files were parsed with `BioPython` v1.81, and per-residue pLDDT values were parsed from the B-factor column. Model confidence in the form of predicted local distance difference test (pLDDT) was used as a proxy for intrinsic-disorder propensity (***Akdel et al., 2022***). Notably, ***Akdel et al. (2022)*** mostly use window-averaged pLDDT profiles; here, we instead use the raw per-residue pLDDT values and apply the same conventional threshold to label residues with pLDDT below 70 as ‘disordered’ and residues with pLDDT above 70 as ‘ordered’. Residue-level secondary structure codes, accessible surface area (ASA), and relative accessible surface area (rASA) were calculated using DSSP from BioPython v.1.81 with default settings (***Rost and Sander, 1994***). Positions with rASA<0.20 were classified as ‘buried’, and those with rASA>0.20 as ‘exposed’. Secondary-structure descriptors were obtained by collapsing the eight DSSP one-letter codes into three biologically meaningful secondary structural groups: ‘helix’ comprising H (canonical *α*-helix, 4-12 hydrogen-bond pattern), G (3_10_ helix), and I (*π* helix); ‘strand’ containing E (extended *β*-strand) and B (isolated *β*-bridge residue); and ‘other/coil’ that bundled T (hydrogen-bonded turn), S (bend), and the dash symbol ‘-’, which denotes residues without an assigned regular secondary structure.

Weighted-contact numbers (WCNs) were used to characterise the local packing density of each residue. The score for residue *i* was computed as

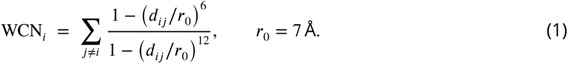

where *d_ij_* is the distance between residues *i* and *j* in the AlphaFold structure and the summation is restricted to residue pairs separated by less than 10 Å. Distances were obtained with MDTraj v1.9.9 using `mdtraj.compute_contacts`. We adopted a ‘side-chain-based’ scheme: representative atoms were the side-chain centroids for standard amino acids and the C*_α_* atom for glycine, which lacks a C*_β_*. We aligned the resulting contact matrix to UniProt residue indices; for proteins whose AlphaFold predictions were delivered as overlapping 1,400-residue segments, per-segment WCNs were averaged across the 1,200-residue overlaps before indexing so that each position contributed exactly one value. The soft weighting function above approaches unity at short range and decays steeply beyond the scale parameter *r*_0_, providing an effective inverse-square distance dependence without imposing a hard cut-off. Residues lacking any neighbour within the 10 Å sphere were assigned WCN = NaN.

For every protein we reduced the residue-level data to a concise set of global descriptors. Continuous metrics—pLDDT, rASA, and WCN—were summarised by their median values across all residues, yielding robust single-number estimates that are insensitive to local outliers. For secondary structure, we computed the percentage of residues falling into each of the three aggregated classes (helix, strand, coil/other) defined above, providing a composition profile rather than a rank statistic. Together, these per-protein summaries constituted the structural feature set used in the concordance and enrichment analyses.

### Population-genetic constraint and clinical data integration

We obtained transcript-level constraint metrics from gnomAD v4.1 (gs://gcp-public-data--gnomad/release/4.1/constraint/gnomad.v4.1.constraint_metrics.tsv); accessed 2024-11-06) from the Google Cloud public bucket using gsutil. The table includes one row per transcript with flags indicating MANE Select and canonical status as well as constraint fields (observed/expected and derived probabilities). From this table we retained the following columns: `gene`, `transcript`, `mane_select` (Boolean), `canonical` (Boolean), `transcript_type`, and the metrics `lof.pLI`, `lof.pRec`, `lof.pNull`, and `mis.z_score`. We filtered to protein-coding transcripts (`transcript_type = protein_coding`) and removed rows with missing or flagged constraint annotations. For gene-level summaries, we selected the MANE Select transcript where available (otherwise the canonical transcript); if neither flag was present for a gene, we chose the transcript with the longest coding sequence. The four metrics were interpreted per gnomAD documentation: `lof.pLI` (probability of loss-of-function intolerance; haploinsufficiency-like), `lof.pRec` (probability of recessive model), `lof.pNull` (probability of being unconstrained), and `mis.z_score` (missense constraint Z-score; higher is more constrained). We note that the v4.x constraint metrics have been iterated since v2 and remain under active refinement by the gnomAD team.

We obtained gene-level estimates of the heterozygous selection coefficient (*S*_het_) from ***Zeng et al. (2024)***, using their public Zenodo repository (https://doi.org/10.5281/zenodo.10403680; accessed 2024-11-12). Specifically, we used version 3 of the file s_het_estimates.genebayes.tsv, which contains posterior estimates from the GeneBayes framework (https://github.com/tkzeng/GeneBayes) alongside auxiliary counts. For downstream analyses we used `post_mean` as the point estimate of *S*_het_ for each gene. We mapped Ensembl gene IDs to UniProt protein annotations using pybiomart (https://github.com/jrderuiter/pybiomart). Genes lacking a reliable Ensembl mapping were excluded from *S*_het_ analyses.

Clinical data was extracted from the ClinVar (***Landrum et al., 2018***) variant calling file (accessed 10-Sep-2024) using BCFtools (***Danecek et al., 2021***). These were subsequently mapped to human reference sequences in UniProt (***The UniProt Consortium et al., 2023, 2025***) release 2024_04 with Ensembl VEP 112 (***McLaren et al., 2016***).

### Benchmarking against multiplexed assays of variant effect

We extracted the reference panel of 36 high-quality multiplexed assays of variant effect (MAVEs) from ***Livesey and Marsh (2025)***. We then benchmarked the selected ten VEPs—`CPT-1`, `AlphaMissense`, `ESCOTT`, `popEVE`, `GEMME`, `VARITY_R`, `ESM-1v`, `SNPred`, `MetaRNN`, and `gMVP`—against these MAVE data using a per-protein, correlation-based procedure. Starting from a variant-level table that included, for each variant, its UniProt accession (`UniProt ID`), one or more MAVE readout columns, and columns for the ten VEP model scores, we grouped rows by UniProt ID and analysed each protein independently.

Prior to benchmarking, we polarity-aligned the predictor scores using global polarity assignments so that larger values consistently correspond to more deleterious variants. For each protein, we first identified the available MAVE readout columns and selected a single representative readout. When multiple candidate MAVE columns were present for a protein, we computed Spearman’s rank correlation (*ρ_S_*; `scipy.stats.spearmanr`, omitting NaN values) between the MAVE values and each available VEP model column using pairwise-complete observations restricted to variants with non-missing values in the MAVE and the corresponding VEP (i.e., a MAVE-anchored, per-model variant set). To ensure robustness to residual score orientation differences and scaling effects, we used the absolute value of the correlation, *ρ_S_*. We summarised the set of per-model *ρ_S_* values for each MAVE column by the median and chose as the representative MAVE readout the MAVE column with the largest median absolute correlation across the ten models. If only a single valid MAVE column was available for a protein, that column was used directly without comparison. Proteins for which no valid MAVE-VEP pairwise correlation could be computed (e.g., due to missing data) were flagged as having no valid MAVE data and excluded from downstream summaries.

Using the selected MAVE readout for a protein, we then re-estimated per-model performance by recomputing *ρ_S_* between that MAVE column and each VEP model column available for the protein. All correlations at this stage were computed on a single, per-protein, strongly anchored shared variant set comprised of variants with non-missing values for the selected MAVE and all included VEP models, ensuring identical sample sizes for VEP-MAVE and inter-VEP comparisons. We retained the full list of model-correlation pairs (sorted in descending order of *ρ_S_*) and also recorded the median of these absolute correlations as a per-protein summary statistic. Results were stored in a per-protein record containing the chosen MAVE column name (`best_MAVE`), the list of `(model,` *ρ_S_* ) pairs (`vep_correlations`), and the median across models (`median_corr`); proteins lacking a valid MAVE readout were stored with `best_MAVE = None`, an empty `vep_correlations` list, and `median_corr = None`. This procedure uses only variants with non-missing values in both the MAVE and VEP columns and applies no additional scaling or normalization beyond polarity alignment of predictor scores, the use of absolute Spearman correlations, and the per-protein selection of a single MAVE assay.

## Code and Data Availability

The data used and produced in this study are available at the Electronic Research Data Archive at University of Copenhagen (KU/UCPH) (ERDA) and available via https://sid.erda.dk/sharelink/glE8N2czUM. The code used to generate the main text figures, to recreate the predictions, and generate new predictions will be made available at GitHub via https://github.com/KULL-Centre/_2026_jonsson_veps.

## Acknowledgments

We thank Benjamin Livesey for help and guidance with collecting the VEP scores and for valuable discussions during the project, and the Atlas of Variant Effects Alliance ‘Analysis, Modelling and Prediction’ workstream for feedback. The research was supported by the PRISM (Protein Interactions and Stability in Medicine and Genomics) centre funded by the Novo Nordisk Foundation (NNF18OC0033950, to KL-L), and by the Medical Research Council (MRC) Human Genetics Unit core grant (MC_UU_00035/9 to JAM). We acknowledge access to computational resources via a grant from the Carlsberg Foundation (CF21-0392).

## Author Contributions

Conception and design: NFJ, JAM and KL-L. Collection of data: NFJ. Analysis and interpretation of data: NFJ, JAM, and KL-L. Reformatting and data availability: NFJ. Supervision: JAM and KL-L. Manuscript preparation: NFJ, JAM, and KL-L.

## Supplementary Material

### Supplementary Tables and Figures

**Table S1.**
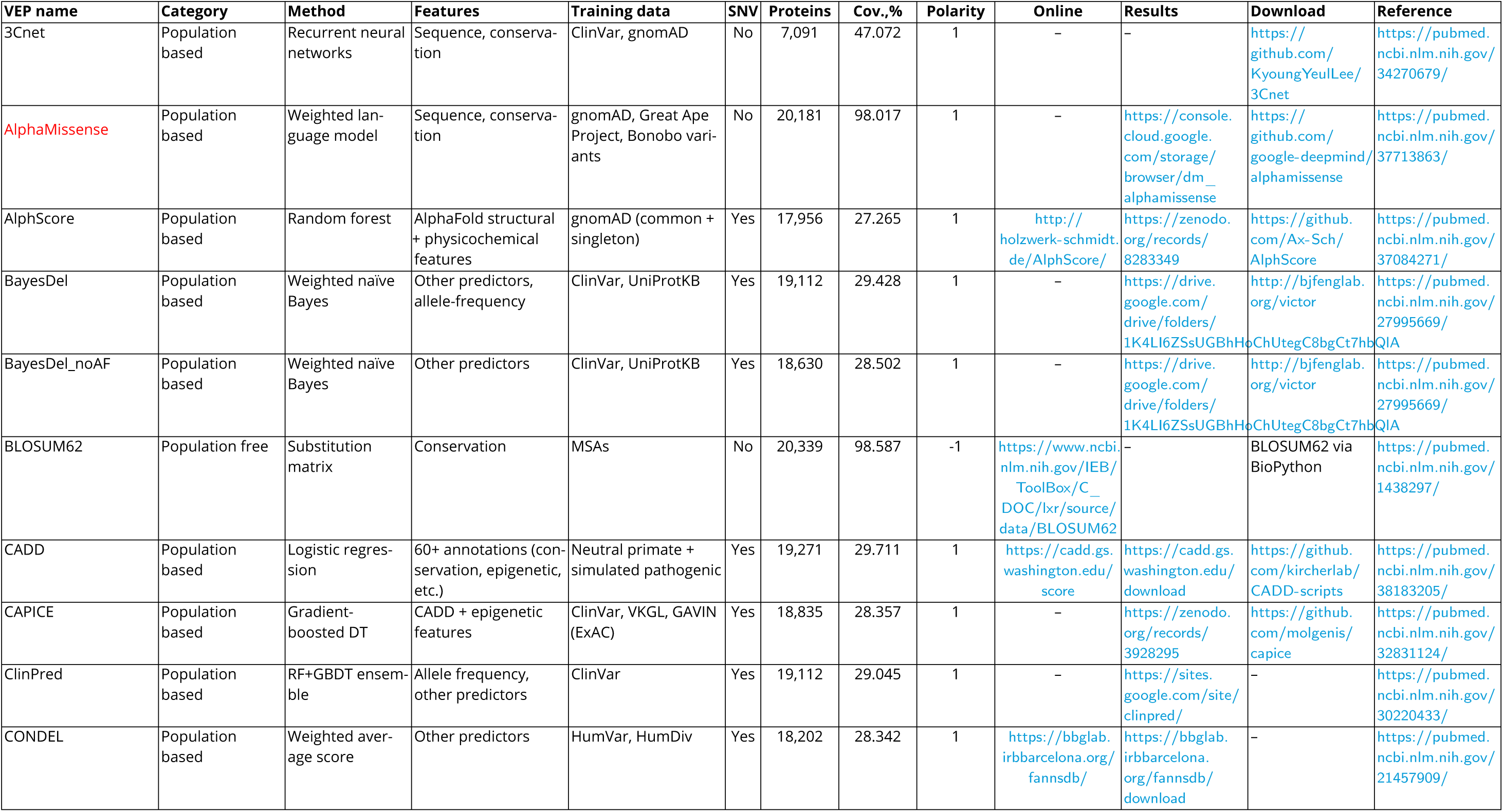
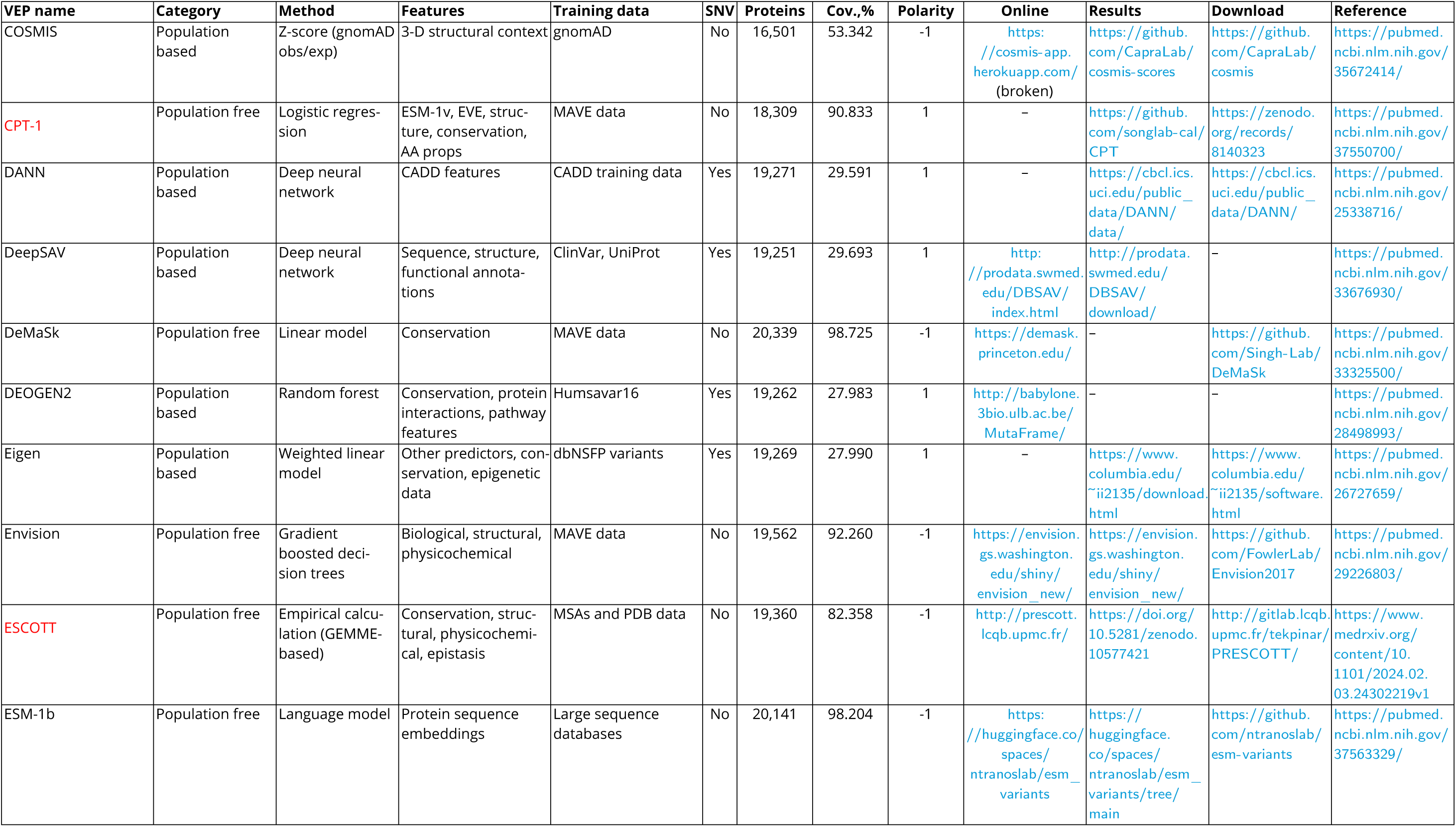
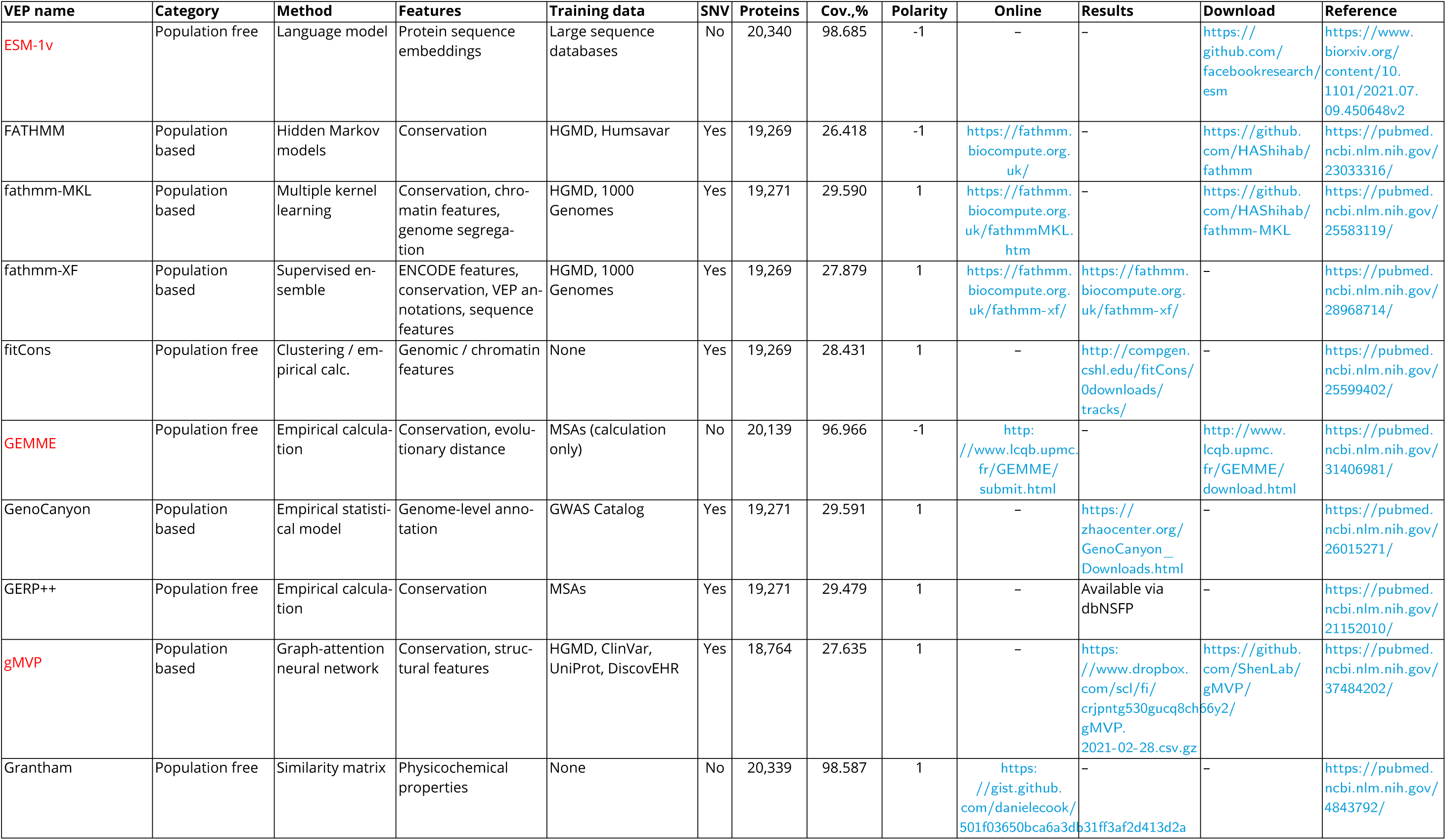
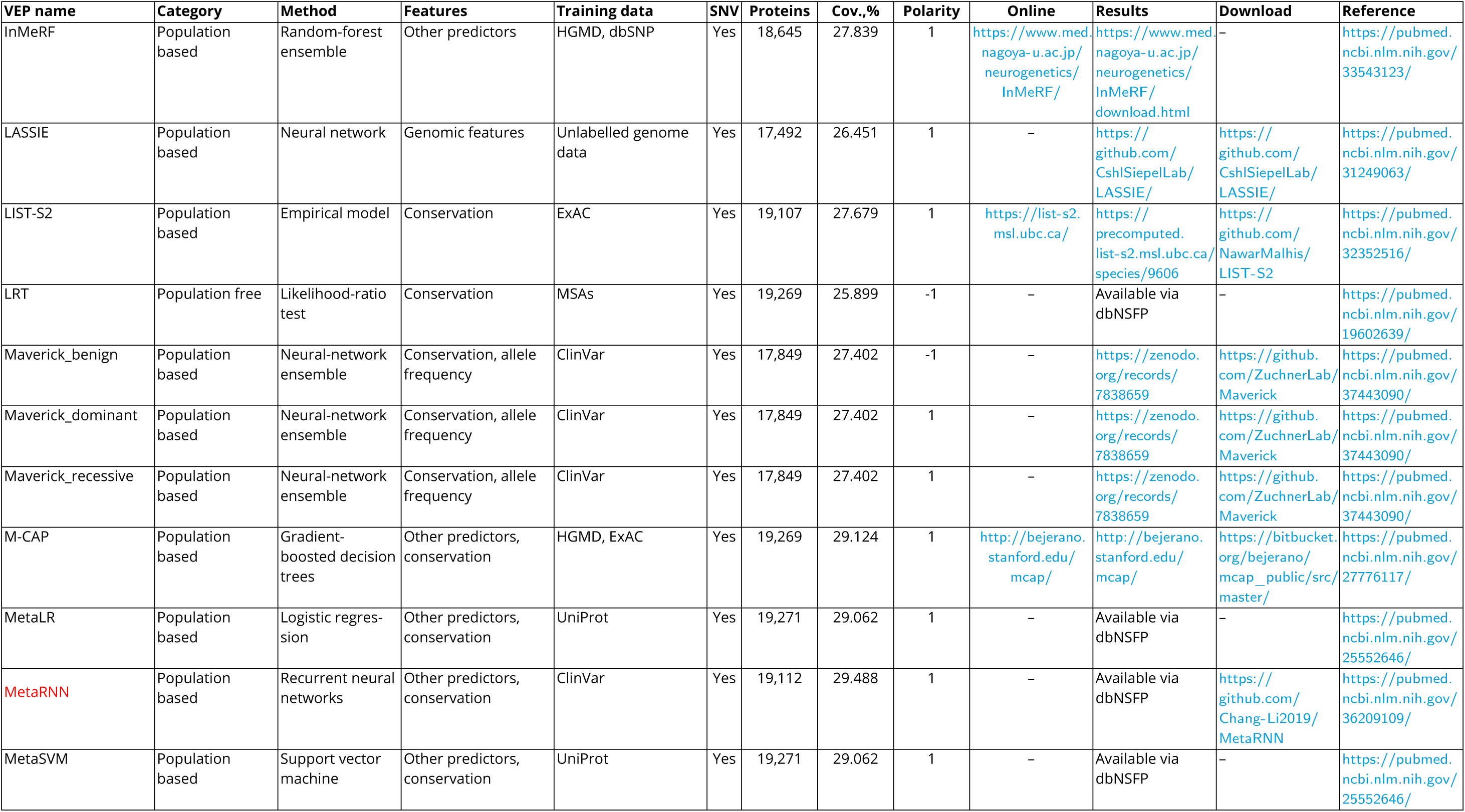
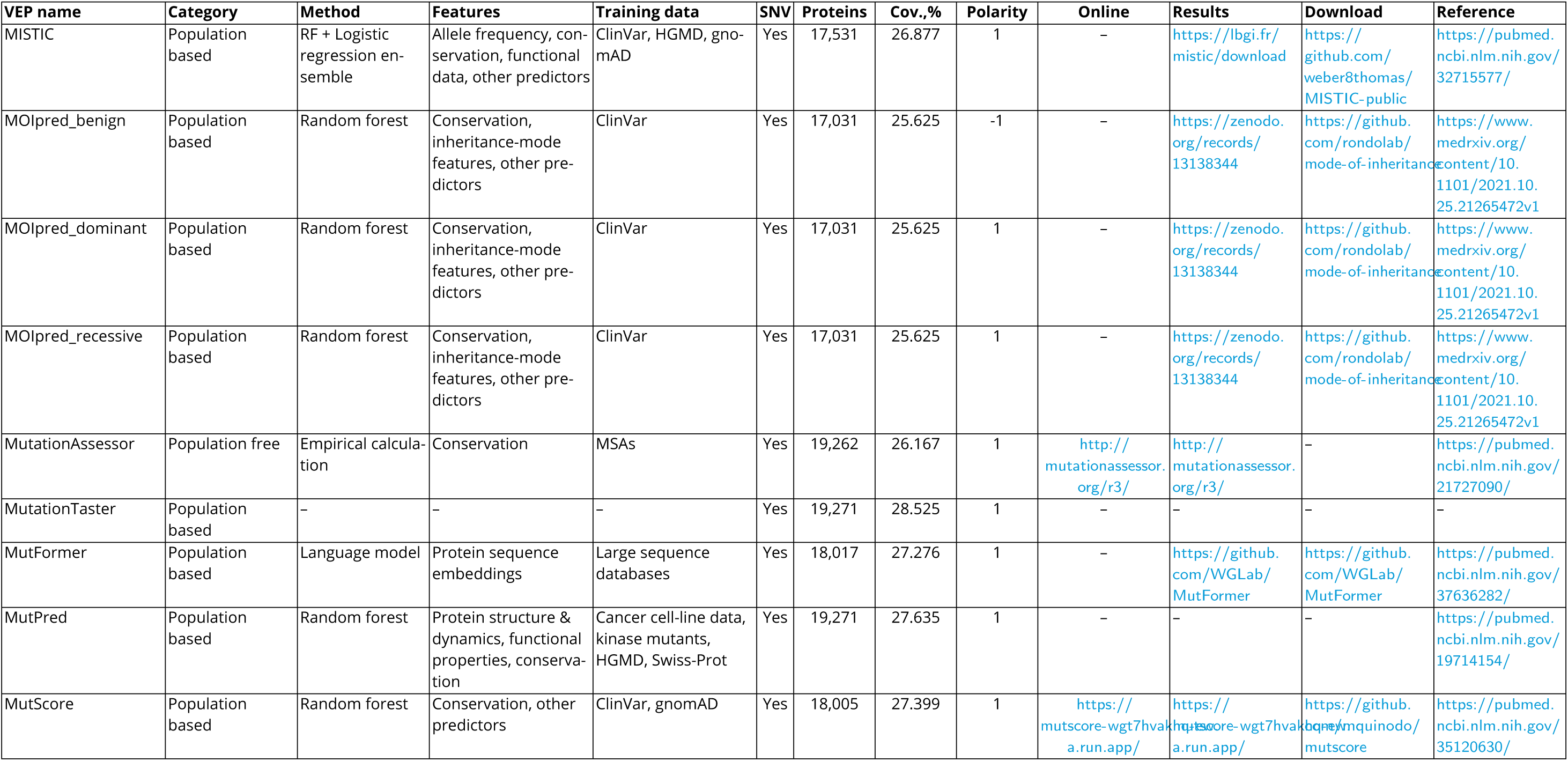
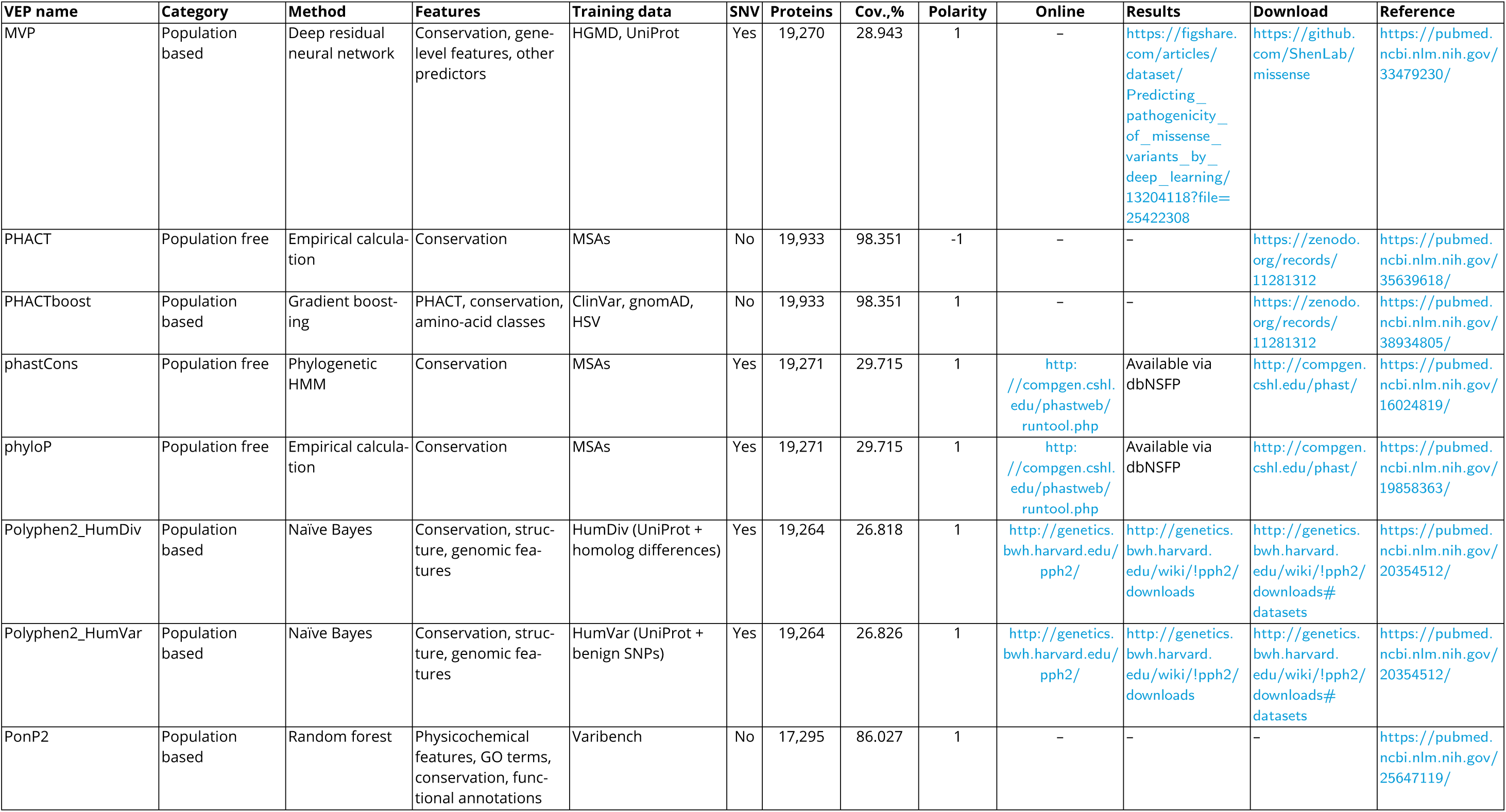
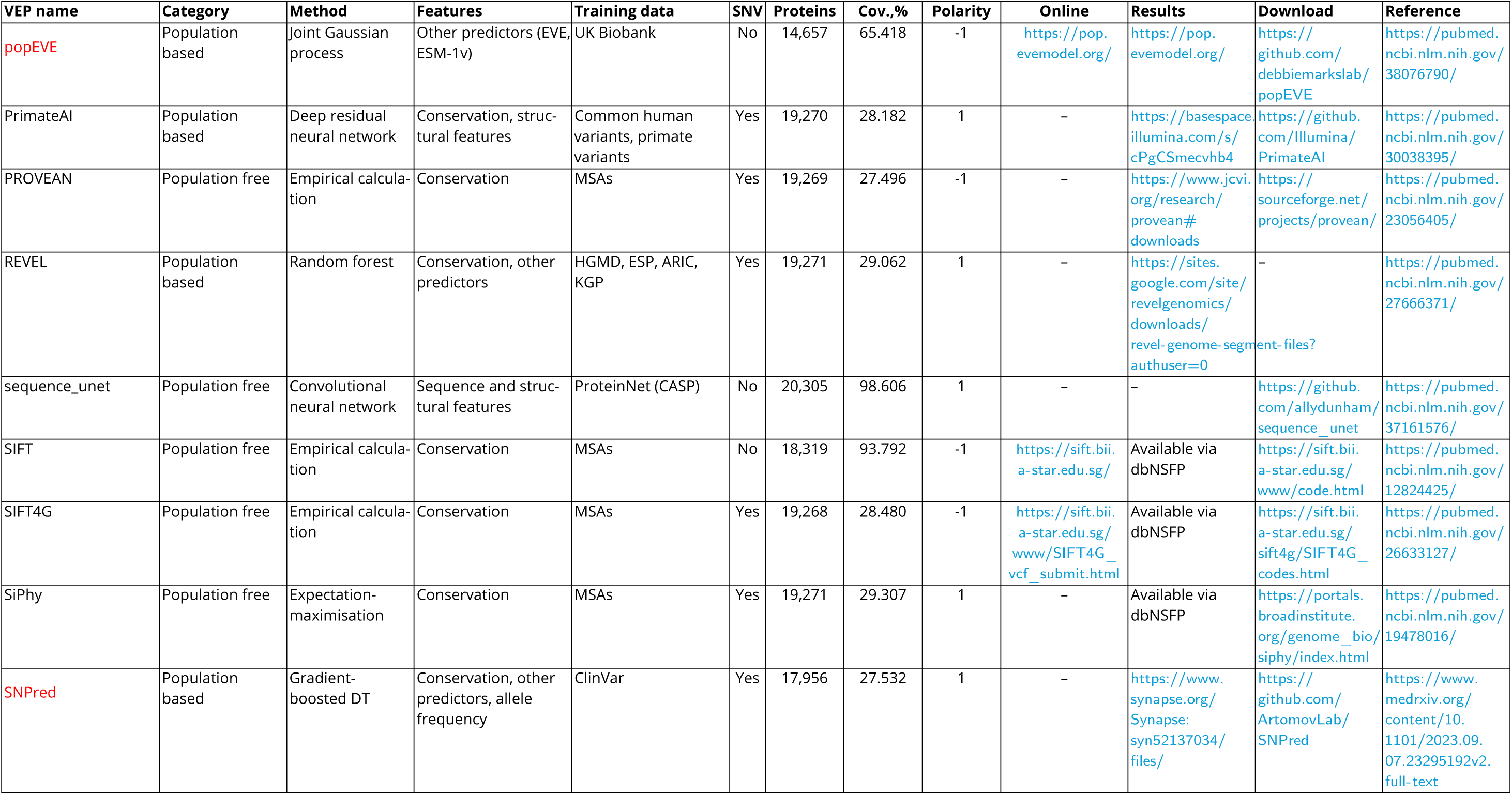
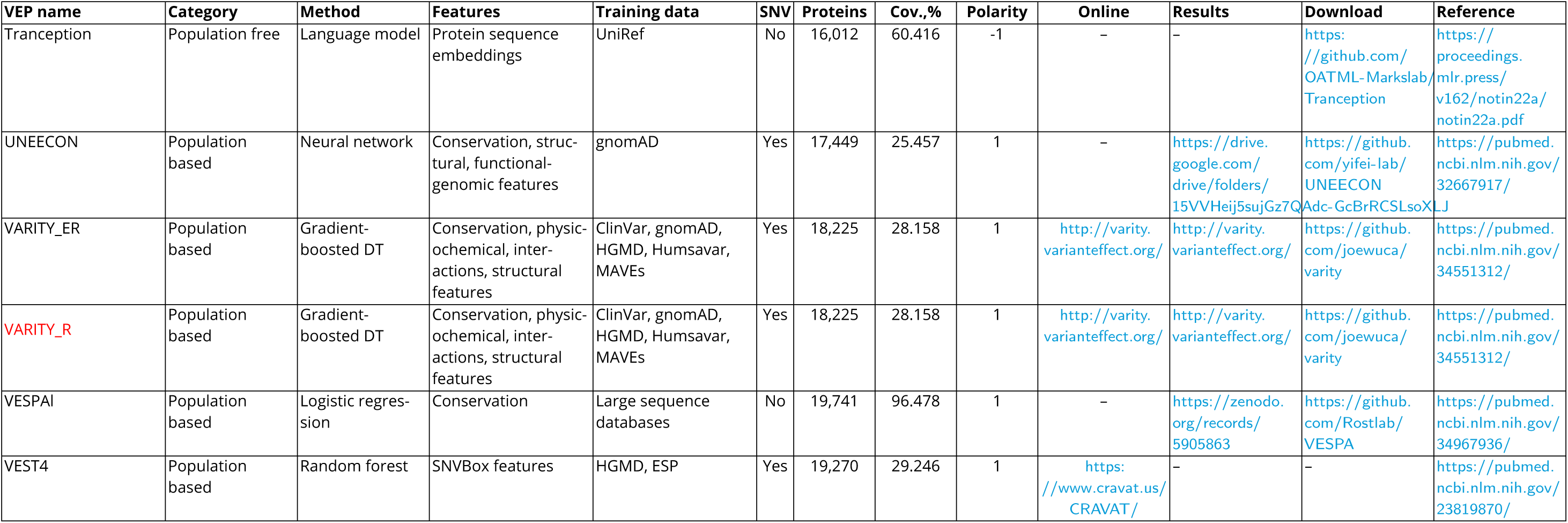
Information on the VEPs in the paper. Columns list the model category according to Livesey et al. and Pathak et al. (***Pathak et al., 2024***; ***Livesey and Marsh, 2025***), learning method, input features, training data, whether the method is SNV-only, protein and residue-coverage metrics, polarity, and resource links. The name of the 10 selected VEPs are coloured with red.

**Figure S1.**
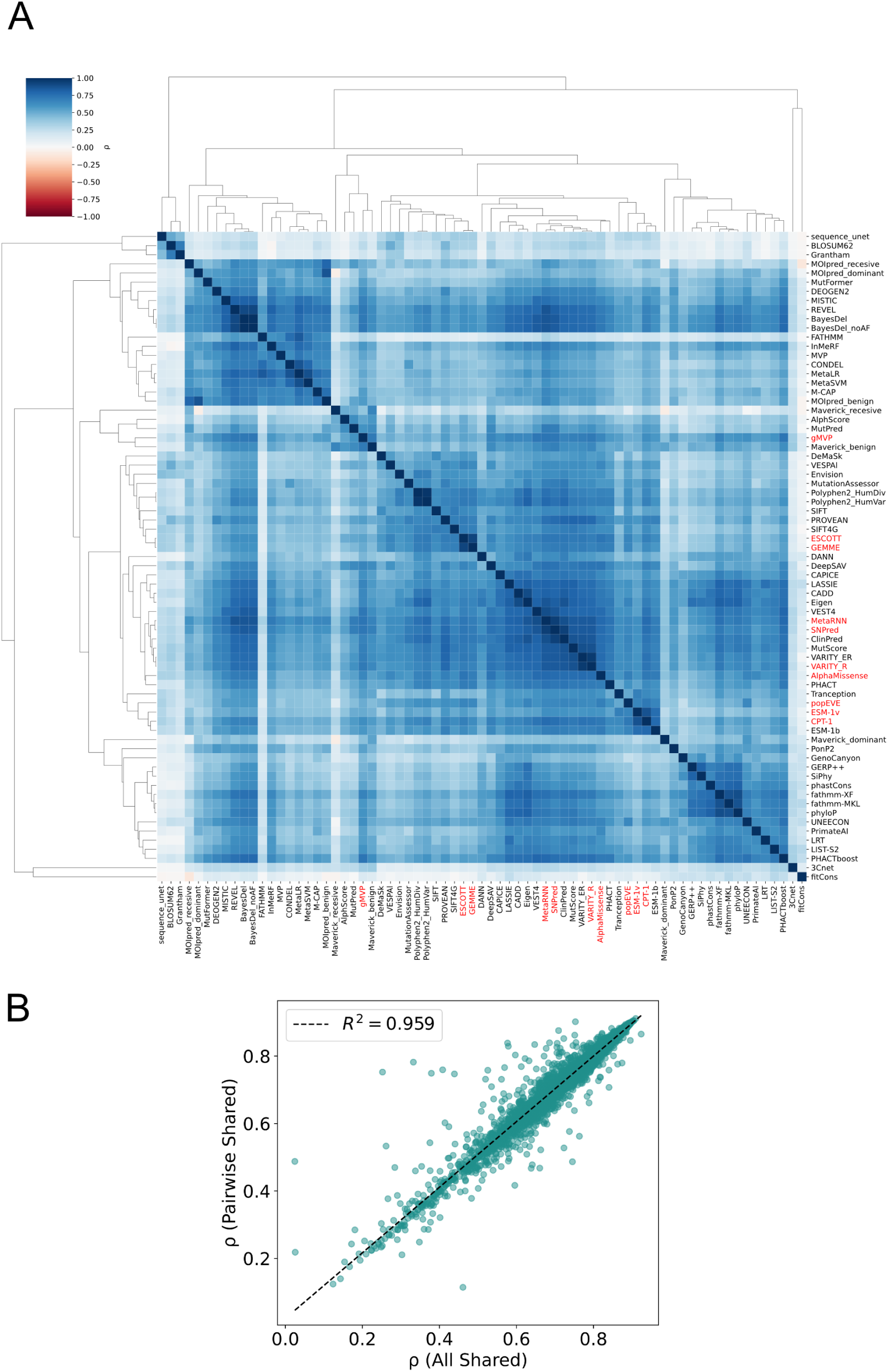
(A) Global concordance landscape across 71 VEPs. The heatmap shows pairwise Spearman correlation coefficients between predictors computed using, for each VEP pair, the maximally overlapping set of variants (including single- and multi-nucleotide substitutions where available) to maximise the number of variants contributing to each comparison. Predictor outputs were first polarity-aligned so that higher scores consistently indicate greater predicted deleteriousness. Rows and columns are ordered by hierarchical clustering of the correlation profiles using correlation distance and average linkage, grouping predictors with similar patterns of agreement across the full panel of methods. Distinct blocks correspond to families of related predictors, including embedding and protein language-model approaches (e.g., AlphaMissense, CPT-1, popEVE, ESM-1b/1v), alignment-driven methods (GEMME, ESCOTT), and clinically trained or ensemble predictors (MetaRNN, SNPred, VARITY_R). The ten predictors used in downstream analyses are highlighted in red on the axes. The colour scale represents Spearman correlation coefficients, where positive values indicate concordant variant ranking between predictors, values near zero indicate little rank agreement, and negative values indicate opposing ranking tendencies. Using maximally overlapping variant sets for each predictor pair yields the same qualitative concordance structure as the all-shared-SNV analysis (Fig. 2). (B) Relationship between correlations estimated from the all-shared-SNV dataset and those estimated using pairwise-overlapping variant sets. Recomputing correlations on the strict intersection of SNVs scored by all 71 VEPs yields highly similar relationships between predictors. Although effect sizes are slightly compressed due to the smaller shared variant set, the overall concordance structure and the relative placement of the ten highlighted predictors remain stable.

**Figure S2.**
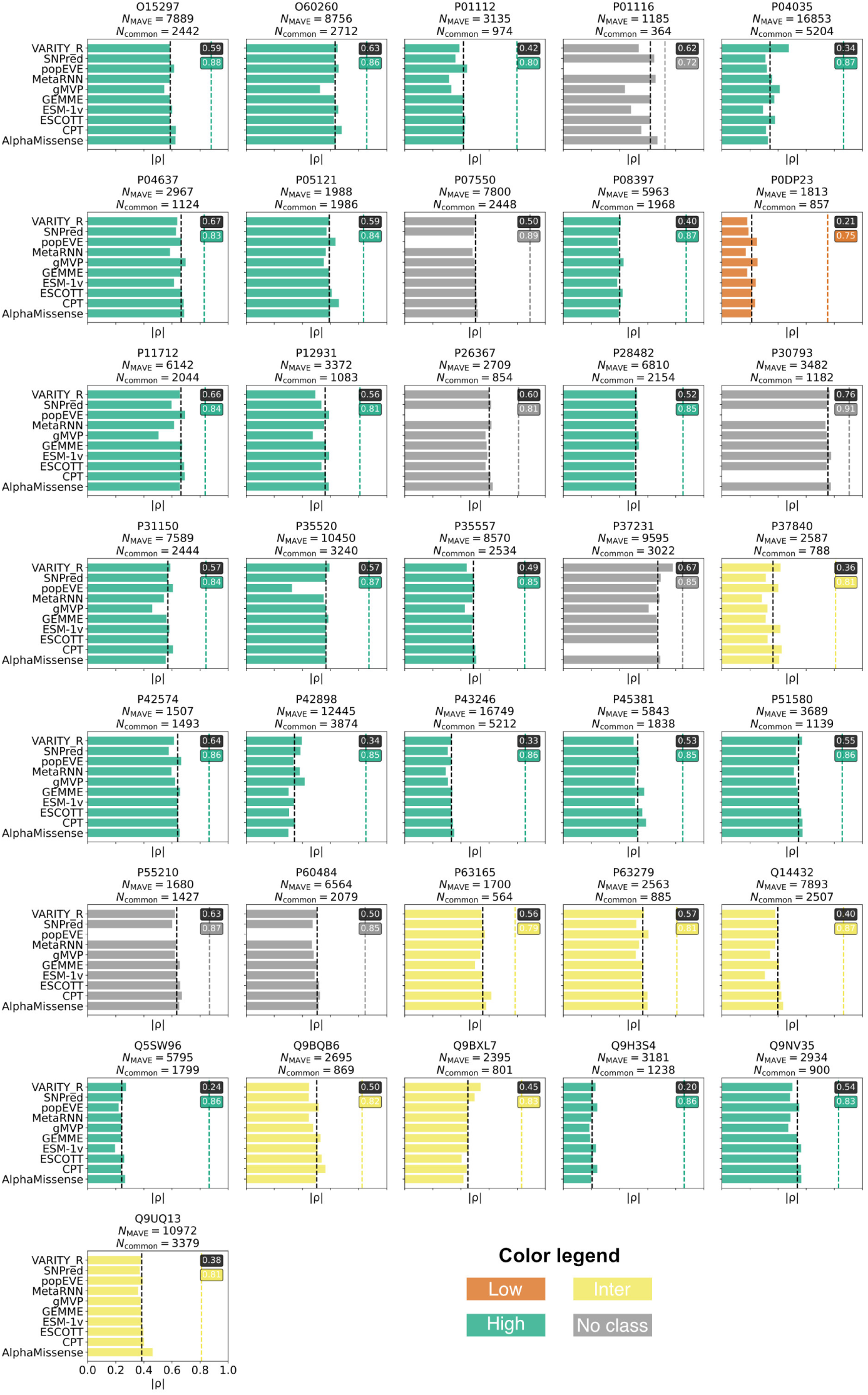
Per-protein VEP-MAVE correlations. Each panel corresponds to one protein with data generated by a MAVE and shows one bar per predictor: the absolute Spearman correlation between that VEP and the MAVE readout, computed on substitutions shared by the assay and the predictors. Two dashed vertical lines summarise each panel: the black line marks the within-protein median absolute Spearman correlation across all VEP-MAVE pairs (experimental predictability; this provides the *x*-axis value in Fig. 6), and the coloured line marks inter-VEP concordance (the median absolute pairwise Spearman among the ten predictors computed on the shared-SNV set; this provides the *y*-axis value in Fig. 6). Panel colouring reflects the inter-VEP agreement tiers defined in Fig. 2 (low, intermediate, high); proteins lacking scores from all ten predictors are unassigned and shown in grey. The grid highlights substantial within-protein spread across predictors and reinforces the absence of a simple relationship between predictor concordance and MAVE alignment; Fig. 6 provides the condensed view of these two summaries.

